# Stability criteria for the consumption and exchange of essential resources

**DOI:** 10.1101/2021.11.24.469922

**Authors:** Theo Gibbs, Yifan Zhang, Zachary R. Miller, James P. O’Dwyer

**Affiliations:** Lewis-Sigler Institute for Integrative Genomics, Princeton University, Princeton, New Jersey, USA; Department of Plant Biology, University of Illinois, Urbana, Illinois, USA; Department of Ecology & Evolution, University of Chicago, Chicago, Illinois, USA

## Abstract

Models of pairwise interactions have informed our understanding of when ecological communities will have stable equilibria. However, these models do not explicitly include the effect of the resource environment, which has the potential to refine or modify our understanding of when a group of interacting species will coexist. Recent consumer-resource models incorporating the exchange of resources alongside competition exemplify this: such models can lead to either stable or unstable equilibria, depending on the resource supply. On the other hand, these recent models focus on a simplified version of microbial metabolism where the depletion of resources always leads to consumer growth. Here, we model an arbitrarily large system of consumers governed by Liebig’s law, where species require and deplete multiple resources, but each consumer’s growth rate is only limited by a single one of these multiple resources. Consumed resources that do not lead to growth are leaked back into the environment, thereby tying the mismatch between depletion and growth to cross-feeding. For this set of dynamics, we show that feasible equilibria can be either stable or unstable, once again depending on the resource environment. We identify special consumption and production networks which protect the community from instability when resources are scarce. Using simulations, we demonstrate that the qualitative stability patterns we derive analytically apply to a broader class of network structures and resource inflow profiles, including cases in which species coexist on only one externally supplied resource. Our stability criteria bear some resemblance to classic stability results for pairwise interactions, but also demonstrate how environmental context can shape coexistence patterns when ecological mechanism is modeled directly.

**Author summary:** One of the longstanding challenges in community ecology is to understand how diverse ecosystems assemble and stably persist. Microbial communities are a particularly acute example of this open problem, because thousands of different bacterial species can coexist in the same environment. Interactions between bacteria are of central importance across a wide variety of systems, from the dynamics of the human gut microbiome to the functioning of industrial bioreactors. As a result, a predictive understanding of which microbes can coexist together, and how they do it, will have far-reaching applications. In this paper, we incorporate a more realistic understanding of microbial metabolism into classic mathematical models of consumer-resource dynamics. In our model, bacteria deplete multiple abiotic nutrients but only grow on one of these resources. In addition, they recycle some of the nutrients they consume back into the environment as new resources. We analytically derive criteria which, if satisfied, guarantee that any number of microbes will coexist. We find that there are special types of interaction networks which remain stable even when resources are scarce. Our theory can be used in conjunction with experimentally determined interaction networks to predict which species assemblages are likely to stably coexist in a specified resource environment.

## Introduction

Pairwise interaction models have informed our understanding of when competitive interactions will lead to stable equilibria. For example, these classic models imply the coexistence of two competing species when the strength of interspecific competition is less than the strength of intraspecific competition, as well as more general stability criteria for large, multi-species systems with randomly distributed competitive interaction strengths [1–3]. On the other hand, models of pairwise interaction strengths do not explicitly model the effect of environmental context, and this context has the potential to refine or modify our understanding of when a group of interacting species will coexist. For example, one species may outcompete another if both compete for and rely on a given resource, but the same two species coexist if that resource is replaced by two alternative resources, each of which is consumed by only one of the two species.

Recent consumer-resource models incorporating the exchange of resources alongside competition have shed light on stable coexistence in systems where interactions are mediated by abiotic resources [4–7]. In these open systems, the environmental context is specified by resource inflow from outside, and stability turns out to depend both on the structure of which species consume and produce specific resources, and on these resource inflow rates. However, this recent theory has focused on a simplified version of microbial metabolism where the depletion of resources always leads to consumer growth. Specifically, the models in [4] assumed either that the impact of consumers *on* resources was proportional to the growth rate resulting from consumption for substitutable resources (ie. the impact of resources on consumers), or else a generalization of this type of assumption [5, 8, 9] for multiplicative colimitation by essential resources.

In general, the rates at which a consumer depletes resources are not always proportional to the benefits that it derives from their consumption. This kind of mismatch between impact and growth can arise for many reasons. One example is ‘waste’, by consumers with large uptake rates [10], where usable resources are degraded and made unavailable for other consumers. An even more basic origin for this mismatch arises when internal metabolism requires multiple resources, but is only limited by the availability of a single resource—known as Liebig’s law [11–14]. In this case, a consumer will deplete multiple resources, but its growth rate will only be sensitive to one of those resources at a time. Thus, a given consumer can strongly affect the growth rate of others that *are* limited by one of its own non-limiting resources.

The consumption of non-limiting resources does not result in cell division. These non-limiting resources could instead be used for cellular maintenance, or transformed via cellular metabolism into byproducts and then leaked back into the environment [15, 16]. For example, a recent reconstruction of the metabolic evolution of the marine cyanobacteria *Prochlorococcus* suggests that *Prochlorococcus* is nitrogen limited and leaks organic carbon, forming a mutualism with the heterotrophic bacterium SAR11 [10]. The depletion of non-limiting resources is one way to generate a mismatch between consumption and microbial growth. Because of the conservation of resource biomass, this mismatch gives rise to the production of resource byproducts, and hence the potential for cross-feeding. Both the mismatch between depletion and growth as well as the cross-feeding of nutrients may have important ecological consequences because they shape the resource environment of the competing species. And yet, we do not know whether or not this more realistic picture of microbial metabolism changes our theoretical understanding of coexistence in diverse microbial communities.

In this paper, we model an arbitrarily large system of consumers undergoing growth governed by Liebig’s law. We consider dynamics near a positive equilibrium where each consumer is limited by a single, distinct resource, but can potentially deplete additional resources. Each consumer leaks the resources it does not use for growth back into the environment as other nutrients, thereby tying the mismatch between growth and depletion directly to cross-feeding. Although we are primarily modeling microbial communities, the mismatch between growth and depletion is a general ecological phenomenon, and our results apply equally well to non-microbial systems. We find that with certain additional assumptions it is possible to analytically derive sufficient stability criteria which guarantee that a feasible equilibrium is stable. These criteria mirror those found earlier for a different form of positive interactions [4], and show that the structure of consumption and production networks, as well as the environmental context, affect the stability of this equilibrium. Our theory generalizes well-known results for low diversity consumer-resource dynamics [17], but also identifies stabilizing interaction network structures which do not have a clear low-dimensional analog. Using simulations, we show that our stability criteria apply to network structures and parameter regimes which do not satisfy our precise mathematical assumptions, including situations where many microbial species coexist on only one externally supplied resource [18, 19]. As a result, our theory could be used to select species assemblages whose consumption preferences and nutrient production networks permit coexistence in a specified resource environment.

## Materials and methods

### A model of the consumption and cross-feeding of resources

We consider a model with five basic biological processes – resource supply, consumption of resources, consumer growth, consumer mortality and cross-feeding. In Fig. 1(A), we illustrate the flow of resources in the ecosystem using a conceptual diagram. Let 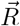 (respectively 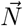) be a vector of *S* resource (respectively consumer) abundances. Resources are externally supplied and depleted through consumption. Let *ρ*_*i*_ be the inflow rate of the *i*-th resource, and let *C*_*ij*_ be the rate of consumption of resource *i* by consumer *j*. We will denote the vector of *ρ*_*i*_ values by 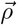 and the *S* × *S* matrix of consumption rates by *C*. We also define *ϵ*_*ij*_ to be the efficiency at which consumer *j* converts resource *i* into new biomass, and we collect these parameters into an *S* × *S* matrix *ϵ* with values in the interval [0, 1]. Each consumer can deplete all the resources in the system, but every consumption event does not lead to new consumer biomass. Instead, we assume that each consumer can only produce new biomass using one resource at a time (which we call the limiting resource), and we define the function 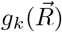 to choose which resource the *k*-th consumer grows on. This is a restrictive assumption, since some bacteria may incorporate biomass from non-limiting resources in nature. At the same time, the contribution to biomass from non-limiting resources is likely to be small compared to the limiting resource, so we are essentially approximating this contribution as zero to preserve analytical tractability.

**Fig 1.**
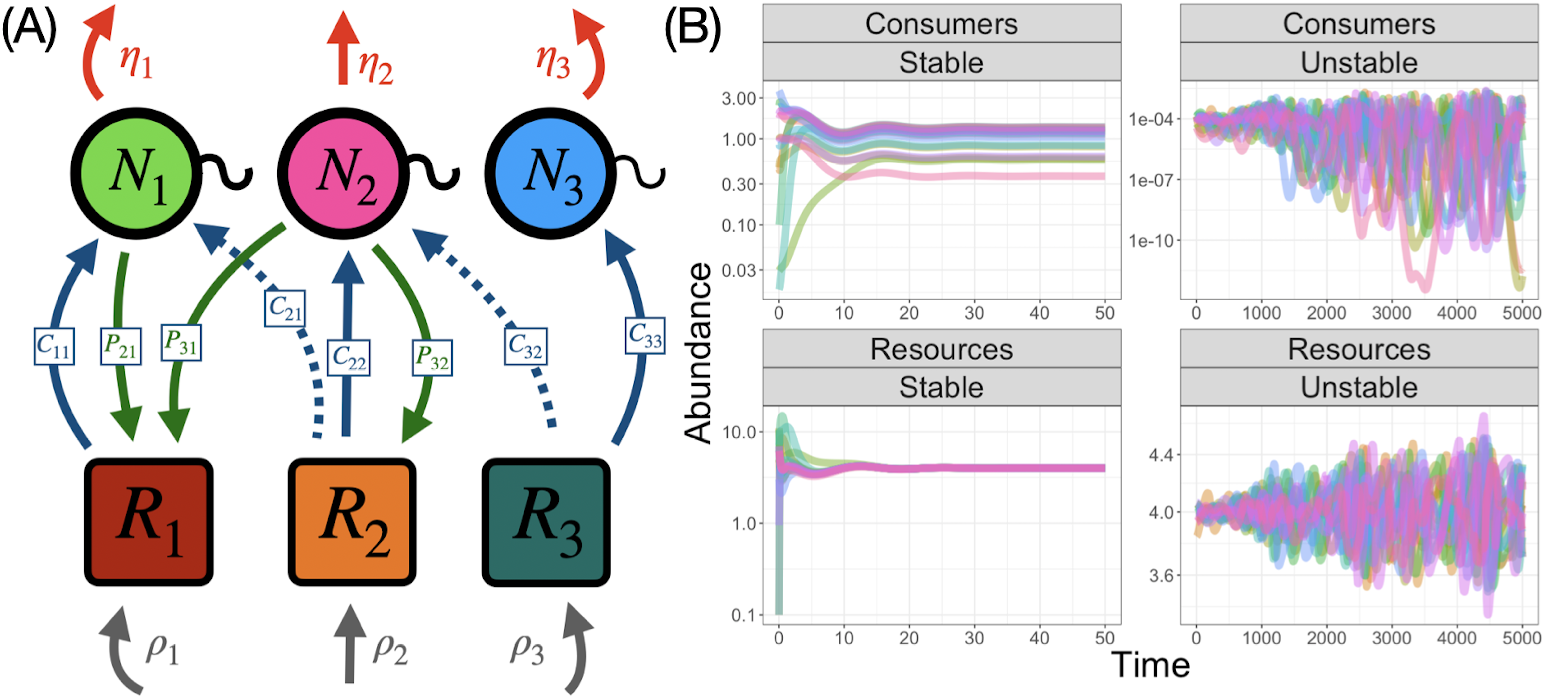
Conceptual diagram and dynamics of consumer-resource model. (A) A schematic of the consumer-resource model in Eq. (1). Consumers (*N*_*i*_) deplete the resource (*R*_*j*_) at rates *C*_*ij*_ (all blue arrows), but only some of the consumption goes to consumer growth (solid blue arrows). Instead, some consumed resource (dashed blue arrows) is leaked back into the environment as new resources at rates *P*_*ji*_ (green arrows). Resources are externally supplied at rates *ρ*_*i*_ (gray arrows) and consumers undergo density independent mortality at rate *η*_*k*_ (red arrows). (B) The dynamics of the consumers and resources in Eq. (1) when there is a stable equilibrium and when there is not. Consumers grow according to the structured Liebig’s law rule described in the text. When there is no stable equilibrium, the consumer abundances undergo large fluctuations, reaching low abundances.

This kind of growth behavior could be produced by a variety of different biological mechanisms. For example, under Liebig’s law, each consumer’s growth is determined only by the resource which contributes the least to the production of new biomass. Therefore, the growth rate of species *k* is given by 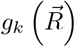 = min_*j*∈{1,…,*S*}_{*ϵ*_*jk*_*C*_*jk*_*R*_*j*_ : *C*_*jk*_ ≠ 0} where we have required that *C*_*jk*_ ≠ 0 so that the minimum is only over the resources which the *k*-th consumer depletes. In nature, microbial species likely cannot grow on every resource they encounter. Therefore, we mainly consider a different version of Liebig’s law (which we call structured Liebig’s law) in which the minimum is only taken over a subset of the resources. Although we focus on these two versions of Liebig’s law, our main results apply equally well to any growth rate function which selects a single resource. For example, if we replace the minimum in the growth function by a maximum, we can interpret the resulting dynamics as a simple model of preferential resource utilization, although the resulting growth behavior differs from the classic understanding of diauxic shifts, because consumers still deplete the resources they are not growing on [20–23].

In addition to growth, consumer *k* undergoes density independent mortality at rate *η*_*k*_. Alternatively, we also set *η*_*k*_ = *η* and interpret it instead as a constant washout rate as in serial dilution experiments [24]. Lastly, consumers recycle nutrients back into the environment by cross-feeding. We define *P*_*ji*_ to be the fraction of consumed resource *j* that is recycled back into the environment as resource *i*. To ensure conservation of biomass, we require that Σ_*i*_ *P*_*ji*_ = 1. As a result, the system as a whole is always competitive, because the overall production of resources is bounded by their total consumption, even though there can also be net flows of one resource to another. For tractability, we assume that each resource can only be secreted as a characteristic set of other resources. Therefore, the resource production rates are the same for every consumer, and *P* is an *S* × *S* matrix, rather than a tensor. We define *δ*_*jk*_ to be an indicator variable that is 1 if consumer *k* is growing on resource *j*, and zero otherwise. Then, 1 − *ϵ*_*jk*_*δ*_*jk*_ is the proportion of resource *j* consumed by consumer *k* that is recycled back into the environment. All together, the dynamics of resources and consumers are given by

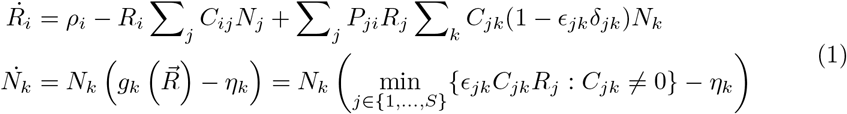

when the consumers grow according to Liebig’s law. In our model, consumers grow on a specific resource, but they can consume and subsequently produce all the resources in the ecosystem. Therefore, each consumer can regulate the resources that the other consumers require to grow. We focus on the equilibria of the model in Eq.1. Specifically, we determine whether or not the small perturbations decay back to the equilibria of Eq. 1 by finding sufficient conditions for the eigenvalues of its Jacobian to have all negative real part. In Fig. 1(B), we plot two different examples of the dynamics of the model in Eq. (1) for consumers following our structured Liebig’s law growth rule. In one example, there is a stable equilibrium to which the consumer and resource abundances converge, while in the other, both consumers and resources undergo large fluctuations. In the following sections, we derive criteria for these equilibria to exist and be stable.

### Equilibria of the consumer-resource model

At equilibrium of the dynamics in Eq.(1), each resource is limiting for precisely one consumer because of the competitive exclusion principle [25–27]. Close to equilibrium, the minimum of the growth rule 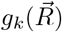 is realized by a unique resource for each consumer. In other words, there is a one-to-one correspondence between consumers and resources at equilibrium. By re-ordering the columns of *C* so that its diagonal corresponds to the limiting consumer-resource pairs, we can rewrite the dynamical system in (1) as

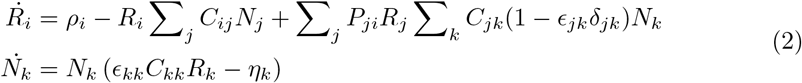

where the *δ*_*jk*_ no longer vary in time. In the S1 Appendix, we plot the dynamics of models in Eq. (1) and Eq. (2). After a transient period in which the dynamics of these two models differ, the dynamics of both models converge to the same equilibrium. More generally, whenever we find an equilibrium of the more complex model in Eq. (1), we can find a corresponding model in the form of Eq. 2 with the same equilibrium properties. Therefore, we restrict our attention to the stability properties of the model in Eq. (2) because this model captures the behavior of models with more biologically realistic growth rules in a neighborhood of equilibrium.

Before we derive stability criteria for the equilibria of the dynamics in Eq. (1), we must determine whether or not there are equilibria of the model in the first place. Specifically, we want to characterize when there are equilibrium solutions where all species coexist at positive abundance (called feasible equilibria [28]). Let’s first analyze the simplified system in Eq. (2). The resource abundances at equilibrium are immediately determined by the consumer dynamics to be 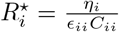. Let’s define 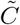 to have the same values as *C* except for the diagonal entries which are given by *C*_*ii*_ (1 − *ϵ*_*ii*_). Then, the equilibrium abundances of the consumers are given by

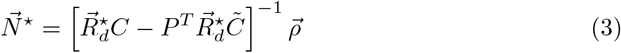

where 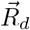 denotes the diagonal matrix with entries given by 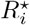. When each of the consumer abundances is greater than zero (*N*_*i*_ > 0), the equilibrium is feasible and it is possible for all species to coexist at abundances that do not change over time. The feasibility of a given set of equilibrium abundances 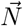 depends on both the consumption and production matrices, as well as the resource inflows and washout rates. In general, it is a difficult problem to characterize the set of feasible abundances in terms of the other parameters of our model. Moreover, depending on the growth rule of Eq. (1), the feasibility of the resource abundances may introduce additional constraints on the interaction patterns. For example, if the consumers grow according to Liebig’s law, corresponding to the minimum growth rule described in the previous section, then there is an upper limit on how large the diagonal coefficients of *C* can be before at least one consumer becomes limited by a different resource at equilibrium. We use a combination of theory and simulation to ensure that we are analyzing the stability of equilibria that are actually part of the dynamics in Eq. (1).

### Sufficient stability criteria for constant abundances

For the sake of analytical tractability, we focus on a simpler version of the model presented in Eq. (1) in this section. Specifically, we take *ϵ*_*ii*_ = *ϵ*, *η*_*i*_ = *η* and we let all the diagonal values of *C* be given by *C*_*d*_ so that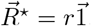. We also assume that 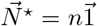 by choosing a corresponding resource inflow vector 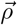. Throughout this paper, we take the diagonal consumption coefficients to be the same for each consumer (*C*_*ii*_ = *C*_*d*_), but we relax the assumption that all the resources (respectively consumers) have equal abundances in later sections. We state two criteria which, if satisfied, imply that the equilibrium where all species coexist is stable.

1. The matrix 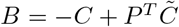 is symmetric.
2. All of the eigenvalues of *B* are negative.

The first criterion measures the reciprocity of the interactions between the consumers in the ecosystem. The matrix element *B*_*ij*_ is the difference between *C*_*ij*_ (which is the consumption of resource *i* by consumer *j*) and 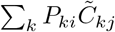 (which is the total production of resource *i* mediated by consumer *j* through the recycling of all other nutrients in the system). Overall, *B*_*ij*_ is the net effect that consumer *j* has on resource *i*. Therefore, if *B* is symmetric, then consumer *j* alters the dynamics of resource *i* in the same way that consumer *i* alters the dynamics of resource *j*. Since each consumer grows on only one resource, we can interpret the symmetry of *B* as perfectly balanced pairwise competition for each limiting resource. Consumer *i* grows only on resource *i*, so consumer *j*’s effect on resource *i* (and hence consumer *i*’s growth) is exactly matched by consumer *i*’s effect on resource *j*. Although we don’t expect natural systems to be precisely symmetric, the symmetry of *B* represents an interesting limiting case of our analysis. Moreover, previous work has found that interaction networks are stabilized by being near to an exactly reciprocal network structure [4, 5, 29]. In the Results and S1 Appendix, we find the same behavior in our model – as the matrix *B* becomes close to a special network structure, it is more easily stabilized.

The second criterion shows that the stability of the matrix *B* informs the stability of the ecosystem as a whole. There are many ways to generate a stable matrix, but one possible method to make *B* stable is to require that its diagonal elements are large compared to its off-diagonal elements. From the Gershgorin circle theorem [30], if

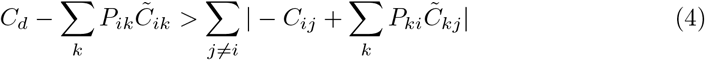

then the matrix *B* is stable, and therefore so is the equilibrium (see the S1 Appendix for a more complete discussion of the Gershgorin circle theorem). We note that this is a sufficient, but not necessary, condition for stability, so the direct stability of the matrix *B* itself yields a more precise understanding of the stability of the equilibrium. Nevertheless, the bound in Eq.4 admits a clearer ecological interpretation. Specifically, satisfying Eq. 4 ensures that each consumer regulates its own resource more strongly than the other consumers affect it, mirroring classic results on consumer-resource models for two resources [17]. Since *B* is symmetric from the first criterion, Eq. 4 is equivalent to requiring each consumer to more strongly alter the dynamics of its resource than it alters the dynamics of all the other resources in the system. The difference between consumption and production of each resource enters the right-hand side of the inequality, so large mismatches between consumption and production are destabilizing, rather than large consumption or production coefficients directly. For example, if there is a resource which is disproportionately used to produce other resources, then the right-hand side of the inequality in Eq.4 will be large and this criteria could be violated. An extreme example of such a destabilizing production matrix *P* would have a column of all ones and zeroes elsewhere. By contrast, if the production matrix is the identity so that there is no recycling of nutrients, then the right-hand side of the inequality is exactly zero and the inequality is always satisfied as long as the consumers can grow at all (*ϵ* > 0). Similarly, if all consumption coefficients and production coefficients are equal, then the right-hand side of the inequality will be relatively small as well.

In summary, the first criterion requires the interactions in the system to have a specific structure, while the second criterion ensures that the strength of the inter-species interactions is not too large relative to the intra-species interactions in the community. The first criterion is not sensitive to the magnitude of the consumption or production coefficients, as long as they are arranged in such a way to produce a symmetric *B* matrix. In the next section, we describe two different matrix parameterizations which generate *B* matrices which always satisfy the first criterion, regardless of whether or not the *B* matrix is stable. Because the second criterion measures the relative strength of the interactions, it does depend on the magnitude of the consumption and production coefficients. Importantly, the parameter *C*_*d*_ controls the intra-specific competition for all the consumers, and it is present on both sides of the inequality in Eq. 4 since it is in the sum 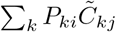. In the following sections, we find that, as *C*_*d*_ increases, the second criteria is more likely to be satisfied. Therefore, we use it as a bifurcation parameter throughout this work by computing the smallest *C*_*d*_ value at which the equilibrium becomes stable.

### Example consumption and production structures

In this section, we describe the three interaction matrix structures which we analyze most closely, but we list all the matrices we tested in the S1 Appendix, as well as brief descriptions of how we generate the matrices. We consider two different methods of parameterizing the consumption matrix *C* and the production matrix *P* that satisfy our first stability criterion, and we display them in Fig. 2. First, let’s consider an ecosystem where each resource is equally re-distributed back into the environment among the other resources. We take 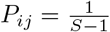 for all *i* ≠ *j* and *P*_*ii*_ = 0, so that each consumer produces an equal mix of all the other resources as it consumes resource *i*, except for resource *i* itself, which it does not produce. We call this case the constant production matrix case. We also assume that *C* is a symmetric matrix with row (and column) sums that are all equal. In particular, let Σ_*j*_ *C*_*ij*_ = *C*_*d*_ + (*S* − 1)*C*_0_ for each *i* so that the average value of the off-diagonal elements of *C* is *C*_0_. We term this parameterization the tradeoff case, since each consumer has a fixed overall consumption strength. We interpret the assumption that each consumer has a fixed total consumption strength as imposing a kind of resource-independent competitive equivalence between consumers, which might emerge from microbes having similar overall metabolic capabilities, as in [31]. In Fig 2(A-B), we plot examples of these two different matrix parameterizations. In the S1 Appendix, we show that the first stability condition is automatically satisfied (ie. that the matrix *B* is symmetric) and we derive an inequality which implies the second stability criterion:

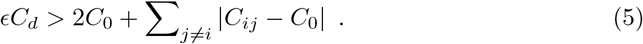

**Fig 2.**
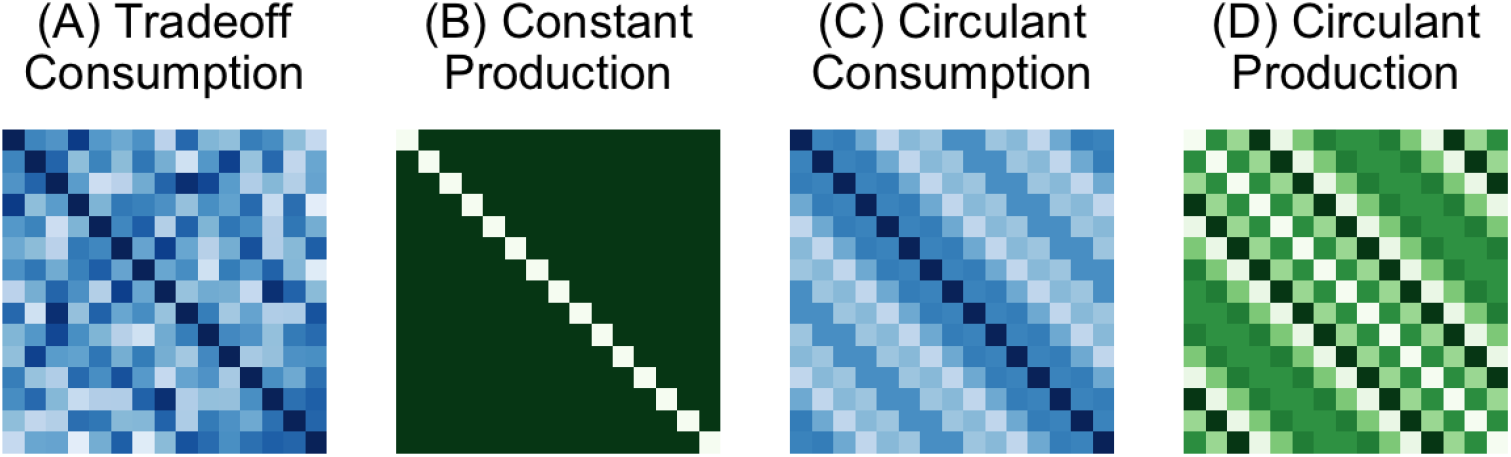
Example consumption and production structures. Each panel is a visualization of the different consumption and production patterns which satisfy our analytical criteria. Darker colors indicate larger values. (A) The tradeoff parameterization of the consumption matrix *C*. *C* is symmetric with identical row (or column) sums and diagonal coefficients all set to *C*_*d*_. (B) The constant parameterization of the production matrix *P*. Each entry off-diagonal entry is set to 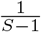, while the diagonal entries are set to 0. (C) The circulant parameterization of the consumption matrix. Each row of the matrix is a permuted version of the previous row and the diagonal entries are all set to *C*_*d*_. (D) The circulant parameterization of the production matrix. Each row of the matrix is a permuted version of the previous row and the diagonal entries are set to 0.

In Eq. (5), the growth rate of each consumer must be larger than twice the average consumption coefficient for all other resources plus the mean deviation of the off-diagonal consumption coefficients. The bound in Eq. (5) has a clear ecological interpretation: as both the mean consumption strength and the variability in consumption strengths increase, consumers must more strongly regulate their preferred resource to maintain stability.

Another class of consumption and production matrices which satisfy our first criterion are symmetric circulant matrices. In a circulant matrix, each row is composed of the same values, but the values are shifted one element right as we descend the rows (see Fig. 2(C-D) for a picture). In this case as well, we set the average value of the off-diagonal elements to be *C*_0_, while the diagonal entries are all exactly *C*_*d*_. If both *C* and *P* are symmetric circulant matrices, then their product is also a symmetric circulant matrix, and therefore the matrix *B* is symmetric as well, so that our first stability criterion is satisfied. Therefore, we need only check the second criteria to determine whether or not ecosystem is stable. As in the constant cross-feeding case, both the average value of the consumption coefficients and their variability affect the stability of the system. We refer to this parameterization as the circulant case.

Having identified two classes of matrices which automatically satisfy our first stability criterion, we also analyze a class of matrices which explicitly violate it. We consider *C* matrices whose off-diagonal elements are sampled from a uniform distribution with a specified mean *C*_0_, while keeping all the diagonal elements of *C* equal to *C*_*d*_. Similarly, we will sample the entries of *P* from a uniform distribution and then enforce the constraint that Σ_*i*_ *P*_*ji*_ = 1. In this fully random parameterization, the matrix 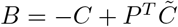 is no longer symmetric, so our first stability criterion is explicitly violated. In contrast, the second criterion can still be satisfied depending on the precise entries in the random matrices *C* and *P*. Therefore, we can test whether or not the first sufficient stability criterion is actually necessary to ensure stability, or if just the second criterion will suffice.

## Results

To evaluate how well our sufficient stability bound predicts the onset of stability in simulated ecosystems, we generated large numbers of random matrices according to each of our three parameterizations for two different values of *C*_0_ and a range of variances. We then numerically computed the smallest *C*_*d*_ value at which the equilibrium becomes stable. In this first set of results, we do not test whether or not the equilibrium is feasible. In the tradeoff and circulant cases, however, we show that feasibility is guaranteed when the consumer (respectively resource) abundances are all equal (see the S1 Appendix for the proof). As a result, only the empirical *C*_*d*_ values from the random parameterizations could be affected by feasibility constraints. In Fig. 3, we plot the average of the numerically-determined *C*_*d*_ values which first induce stability (the points) along with the average value of our sufficient analytical stability bound (solid lines). In each of the three parameterizations, the empirically computed average *C*_*d*_ values are closely predicted by our average analytical *C*_*d*_ values. As a result, the trends in the empirically computed *C*_*d*_ values are reflected by our theory: for both larger mean value *C*_0_ and larger variance in the consumption coefficients, a larger *C*_*d*_ value is required to induce stability (see Fig. 3).

**Fig 3.**
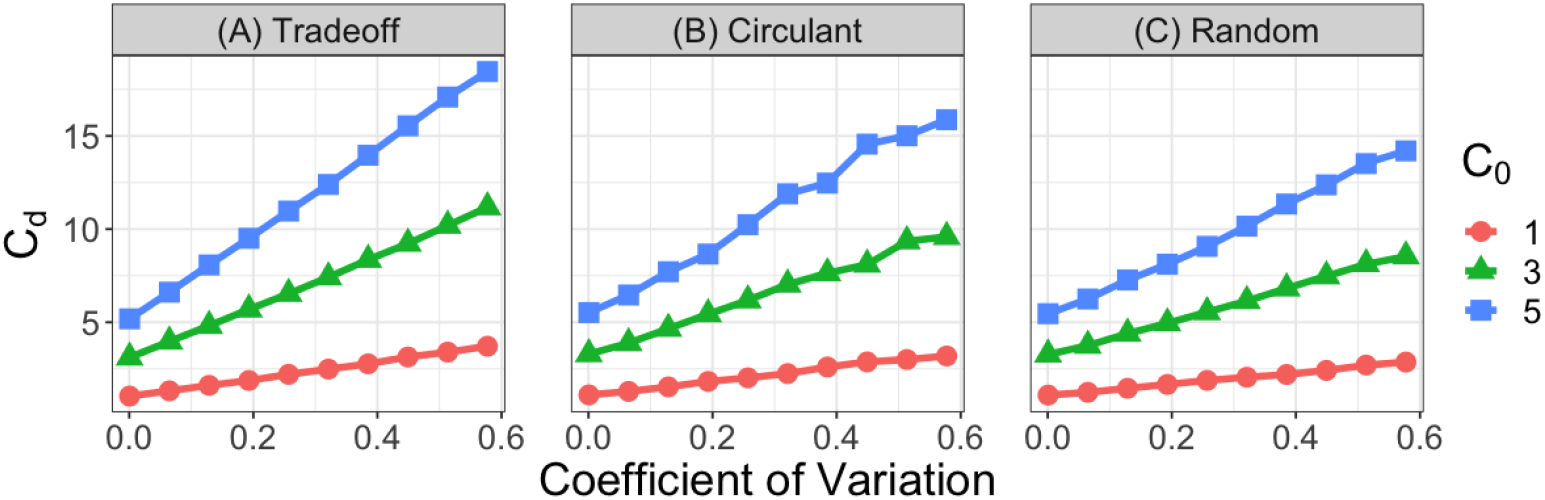
Analytical stability criteria predict empirical values. We plot the value of the theoretical bound for *C*_*d*_ (solid lines) in the second stability criterion averaged over 100 realizations of the random matrices comprising *C* and *P* while varying the standard deviation, and hence the coefficient of variation (the standard deviation divided by the mean), of the off-diagonal elements. The shapes are the smallest *C*_*d*_ value at which the fixed point becomes stable numerically, once again averaged over 100 replicates. The different colors and shapes denote the three different *C*_0_ values, which is the average of the off-diagonal entries. The different panels are for the three different parameterizations of the consumption and production matrices ((A) is the tradeoff parameterization, (B) is the circulant parameterization and (C) is the random parameterization). Parameters: *S* = 15, *n* = *r* = 1 and *ϵ* = 0.5 for all panels. For each of the different matrix parameterizations, we first sample the consumption coefficients from uniform distributions with mean *C*_0_ and the specified standard deviations so that the coefficients of variation vary from 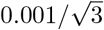 to 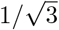. Then, we impose the constraints for the tradeoff and circulant parameterizations afterwards.

In our mathematical theory, we have only proven that our stability criteria are sufficient for, rather than exactly predictive of, stability. And yet, in Fig. 3, our theory accurately predicts the precise average value of *C*_*d*_ at which the system becomes stable. In addition to the average trends of the numerical and analytical *C*_*d*_ values, we also plot the difference between these two quantities for each replicate in the S1 Appendix. For the tradeoff and circulant cases, there is no difference between our predicted *C*_*d*_ values and the numerically determined ones. In fact, in the S1 Appendix, we show analytically that, under specific additional assumptions, the equilibrium is stable if and only if *B* is stable. Our two example consumption and production structures satisfy these assumptions, so our theory exactly predicts when the equilibrium is stable. This combination of mathematical proofs and numerical simulations suggest a more comprehensive result – if *B* is symmetric, then the equilibrium is stable if and only if *B* is stable. This would be a particularly interesting result because it suggests that the simplified matrix *B*, which measures the interactions in the system in an intuitive way, exactly determines the stability properties of the entire community. We cannot prove this statement in its full generality, but in the S1 Appendix, we present additional analytical arguments and numerical evidence in support of it. By contrast, in the random parameterization, the difference between our numerical and theoretical *C*_*d*_ values can be positive or negative, meaning that our analytical criteria do not imply stability. At the same time, our analytical bound predicted the *average* stability properties of the fully random consumption and production matrices remarkably well (Fig. 3(C)), even though our theory is not mathematically justified in this case.

We now more exhaustively simulate our model to identify parameter combinations where our analytical criteria do not predict the average behavior of the empirical *C*_*d*_ values. We numerically compute the smallest *C*_*d*_ value at which the system becomes stable, in addition to our analytical bound, as we vary *r*, *ϵ* and *n*. We find that varying *r* has no effect on the stability or feasibility of the empirically computed *C*_*d*_ values, nor does it affect our analytical bound (see S1 Appendix). We also observe that changing *ϵ* does not affect the empirical *C*_*d*_ values for the tradeoff or circulant cases. Similarly, *ϵ* does not affect the *C*_*d*_ values at which the random case first becomes stable. However, it does affect the empirical *C*_*d*_ values of the random case if we require feasibility as well, because smaller *ϵ* values make it more difficult to find a feasible fixed point (see S1 Appendix).

Most importantly, we find that varying *n* causes the random case to systematically violate our analytical stability criteria. Our theoretical criteria exhibit no dependence on the value of *n*, and neither do the empirically computed *C*_*d*_ values for the tradeoff or circulant cases (see Fig. 4(A-B)). Yet, we find that, as we decrease the consumer abundance *n*, it becomes increasingly more difficult for the equilibrium to be stable for random consumption and production structures (see Fig. 4(C)). Therefore, the specific symmetric structure enforced by our first stability criterion eliminates the effect of *n* on stability, but, for more generic consumption and production patterns, low consumer abundances lead to instability. This result mirrors those of other recent studies, which have also found that special network structures promote stability when consumer abundances are small as a result of low of resource supply [5, 6].

**Fig 4.**
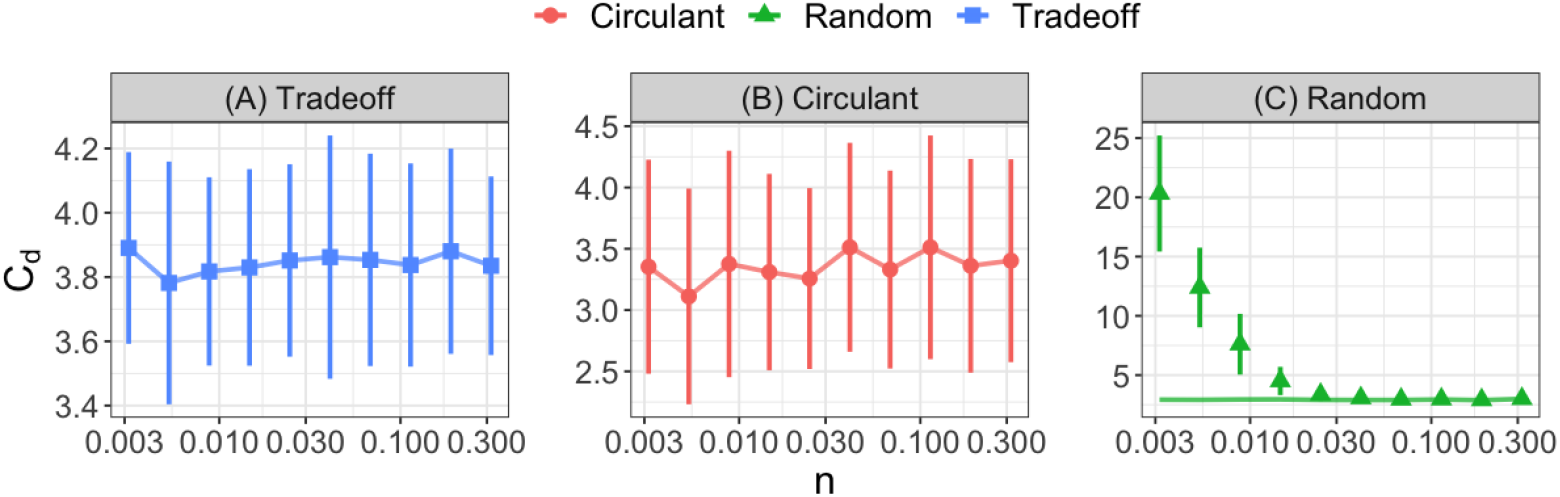
Random consumption structure violates theoretical stability criteria at low consumer abundances. We plot the value of the theoretical stability bound for *C*_*d*_ averaged over 100 replicates (solid lines) as the consumer abundance *n* varies over three orders of magnitude for the three different matrix parameterizations (panels and colors). We also plot the average smallest *C*_*d*_ value for which the system first becomes stable numerically (shapes) along with error bars showing one standard deviation above and below the mean. In panels (A-B), the theoretical bound accurately predicts the dependence of *C*_*d*_ on *n*, while for the random case (panel (C)), the theoretical predictions fail at low consumer abundance. Parameters: *S* = 15, *r* = 1 and *ϵ* = 0.9. The off-diagonal elements of the consumption matrices are sampled from uniform distributions on [0, 2] before the parameterizations are imposed.

In our theory, we assume that the consumer and resource abundances are the same for the different species and resources in the system (ie. 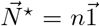 and 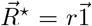). We also treat these abundances as parameters that we can modify. Yet, in experimental microbial communities, 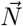 and 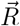 are determined by the resource inflow and dilution rates (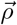 and 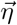), as well as the consumption and production networks. Using simulations, we now ask whether or not the qualitative stability patterns we have derived for the simpler case of constant abundances also describe systems where we choose 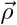 and 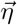 as independent variables and let the dynamics determine the consumer and resource abundances. We set *η*_*i*_ = *η* for all consumers as in a chemostat or a serial dilution experiment. We set *ρ*_1_ = *ρ* and set all other resource supply rates to zero (*ρ*_*i*_ = 0 when *i* = 2, …, *S*), mimicking recent microbial competition experiments [18, 19], where diverse communities coexist via cross-feeding.

In Fig. 5(A-B), we plot the fraction of ecosystems with a feasible and stable fixed point as we vary the total resource inflow *ρ* and the diagonal consumption coefficient *C*_*d*_ for the tradeoff and random matrix parameterizations. We enforce two different aspects of feasibility in these simulations. First, there must be an equilibrium which solves Eq. (3). In other words, there must be a fixed point where all consumers coexist. This constraint means that *C*_*d*_ must exceed a threshold determined by the consumption and production networks (see the lower bound of the coexistence region in the heatmaps in Fig. 5(A-B)). Second, we require that the feasible fixed point found using Eq. (3) also be realized in the dynamics of the structured Liebig’s law growth rule. This second feasibility constraint sets an upper limit on the value of *C*_*d*_. For large enough *C*_*d*_, the minimum of the structured Liebig’s law growth rule is no longer realized by the same consumer-resource pairs, and the feasible fixed point cannot be attained in the time dynamics of the model (see the upper limit of the coexistence region in Fig. 5(A-B)). Both of these feasibility properties, however, are independent of the total resource availability. Instead, decreasing the resource inflow *ρ* leads to lower consumer abundances, but it does not affect whether or not there is an equilibrium with all positive abundances. When the consumption and production matrices are randomly sampled, these low consumer abundances give rise to unstable, but feasible, fixed points once again (Fig. 5(B)). By contrast, the consumption and production networks which obey our first symmetric stability criteria are protected from resource-inflow-mediated instability even in this more complex case (Fig. 5(A)). In the S1 Appendix, we find that these qualitative patterns remain true for other choices of resource inflow rates (ie. randomly sampled 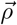 vectors), suggesting that these symmetric consumption and production structures promote stability generally.

**Fig 5.**
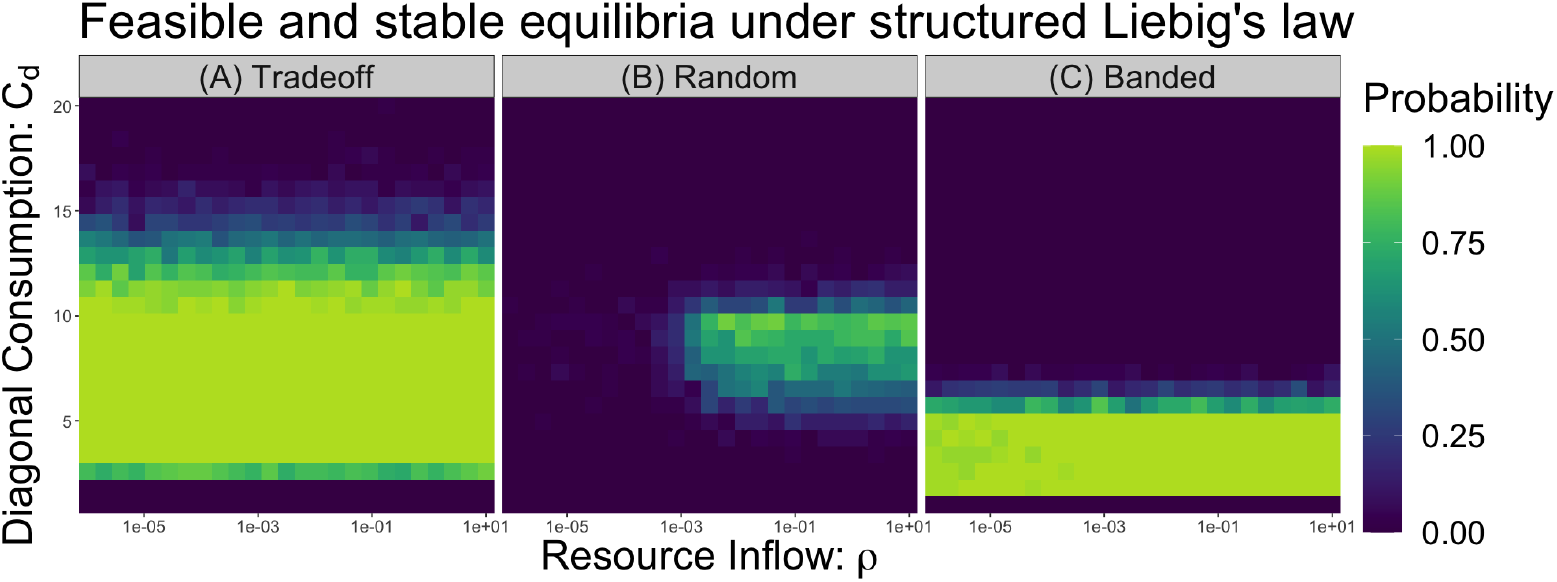
Low consumer abundances induce instability for more general resource inflows and interaction networks. We plot the probability of finding a feasible and stable fixed point in 50 replicates across a range of *C*_*d*_ values and resource inflows *ρ* with only one externally supplied nutrient for three different matrix parameterizations. We also enforce that the fixed point is realized under the structured Liebig’s law dynamics, where each consumer grows on the most limiting nutrient of a subset of the resources. (A) The tradeoff matrix parameterization does not show any dependence on the resource inflow *ρ*, as in Fig. 4. (B) The Random does not have any feasible and stable fixed points at low resource inflow. (C) The Banded matrix parameterization has a consumption matrix with non-zero values on the upper and lower bands of the matrix, displaced from the diagonal by one index. It also has a constant production matrix, as described previously. It does not show any dependence on the resource inflow. Parameters: *S* = 15, *ϵ* = 0.05 and consumption coefficients sampled from uniform distributions on [0.5, 1.5] before the constraints are imposed.

We now construct a variety of consumption and production networks that do not satisfy our symmetric stability condition and ask whether or not they can still protect ecosystems from instability at low consumer abundances. We hypothesize that the symmetric structure of the *B* matrix above protects ecosystems from instability because it ensures that the *B* matrix has purely real eigenvalues. So, we test other consumption and production structures which give rise to non-symmetric *B* matrices with real eigenvalues. In Fig. 5(C), we plot the probability of finding a feasible and stable equilibrium for a case where *C* has non-zero off-diagonal entries only on the upper and lower bands displaced from the diagonal by one index. We constrain the row sums of *C* to be all equal and we take *P* as in the constant case. In this parameterization, *B* is no longer symmetric but it does have real eigenvalues, and we find that the equilibrium remains stable at low consumer abundances 5(C). In the S1 Appendix, we test a suite of other consumption and production structures, and we consistently find the same qualitative results – networks which generate *B* matrices with purely real eigenvalues are stable at low consumer abundances, while *B* matrices whose eigenvalues have non-zero imaginary part are unstable in the same regime. In fact, when there is no cross-feeding (ie. when *P*_*ji*_ = 0 for all *i* and *j*) and the consumption matrix has constant row sums, we prove analytically that, if the consumption matrix *C* has any eigenvalues with non-zero imaginary parts, it is always possible to find parameter combinations where the equilibrium with equal consumer and resource abundances is unstable (see S1 Appendix). Conversely, we show that if *C* has purely real eigenvalues, then the stability of *C* itself implies that the community as a whole is stable.

We don’t expect natural systems to be precisely symmetric or to generate *B* matrices that have eigenvalues whose imaginary parts are exactly zero. At the same time, it is possible that natural systems are better protected from instability if they are near to one of the special interaction structures we have identified analytically. Using simulations, we find that the transition between stability and instability at low consumer abundances is a continuous one. As we increase the correlation between off-diagonal pairs (*C*_*ij*_, *C*_*ji*_) of an otherwise random *C* matrix, the ecosystem can coexist at smaller and smaller resource inflows (see S1 Appendix). Therefore, our results extend to cases in which the *B* matrix does not have precisely real eigenvalues. Instead, the magnitude of the imaginary parts of the eigenvalues of *B* controls the resource inflow levels at which the community can coexist. In short, we conjecture that the matrix *B* serves as a measure of the relevant interactions in the ecosystem, and, when *B* has all real and negative eigenvalues, the equilibrium is always stable. If *B* has some eigenvalues with non-zero, but small, imaginary part, the equilibrium will be stable across a broad range of resource inflows, and the threshold resource supply at which the system first becomes unstable is controlled by the magnitude of the imaginary parts of these eigenvalues.

Our theory predicts when a feasible and stable equilibrium exists in the dynamics of our model, and when such a fixed point does not exist. It does not, however, shed light on what happens to the consumers in an ecosystem without a stable equilibrium. In Fig. 6, we plot the dynamics of two different consumption networks – the banded network and a sparse randomly sampled network – for three different inflow levels of one externally supplied resource. *B* has purely real eigenvalues for the banded consumption network, but some of the eigenvalues of *B* have non-zero imaginary part for the sparse network. Consumers grow according to our structured Liebig’s law growth rule. For the banded network, consumers always converge to a stable equilibrium, while for the sparse network, they converge to equilibrium only when the resource supply is sufficiently large. When the dynamics do not converge to a stable equilibrium, the consumers undergo large, semi-regular fluctuations which vary in amplitude (Fig. 6). In the presence of demographic noise, these large fluctuations would likely lead to exclusion. In addition to these semi-regular fluctuations, consumers can converge to stable limit cycles in some cases. Depending on the initial conditions, it is also possible for some consumers to become limited by a different resource, leading to consumers reaching low abundances or even becoming excluded (see S1 Appendix).

**Fig 6.**
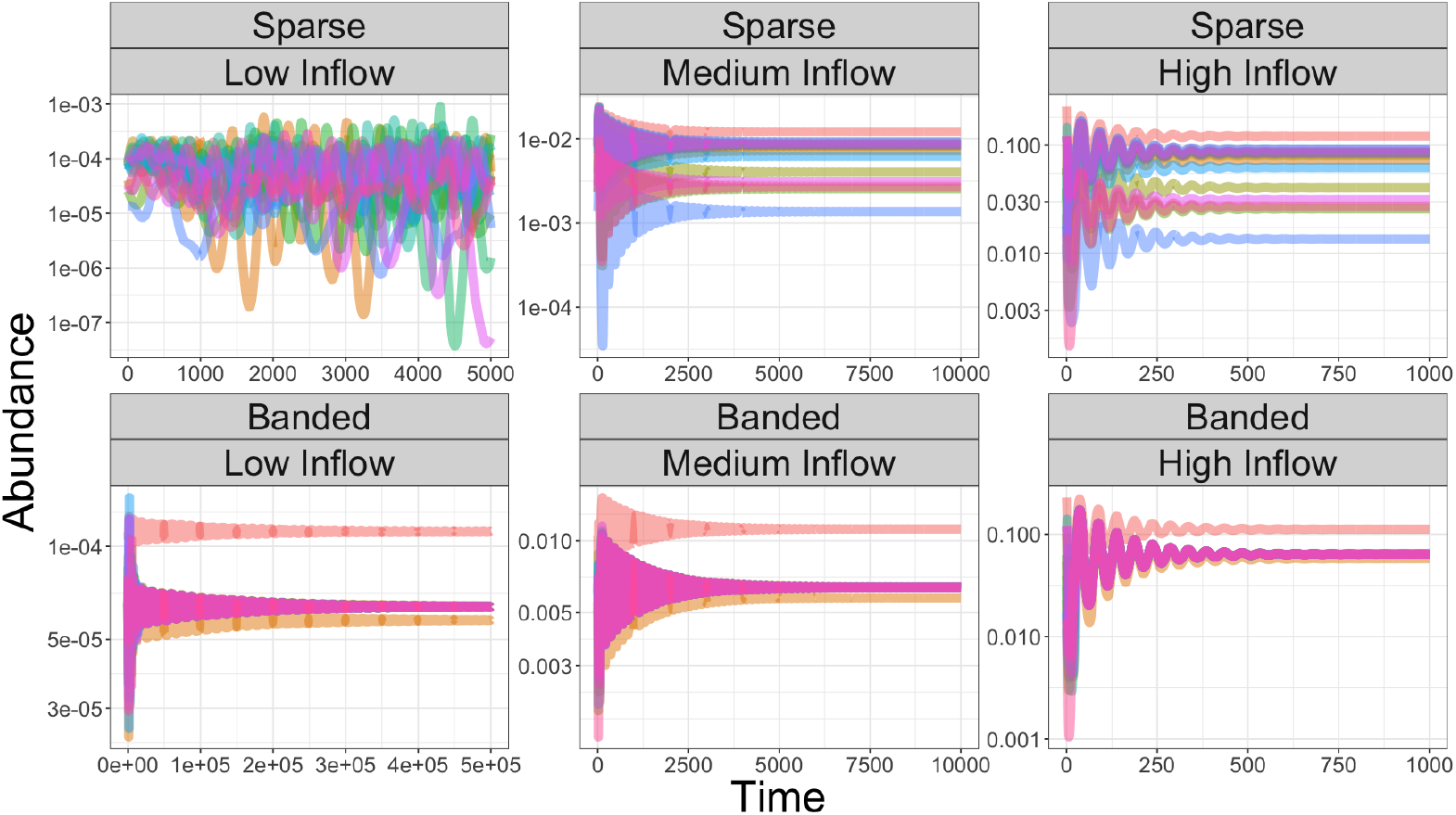
Stable and unstable consumer dynamics for varying resource inflows. We plot the consumer dynamics for two different consumption networks – one where the coefficients are sampled randomly from a zero-inflated distribution (labeled Sparse) and the other according to the banded consumption network described in the text (labeled Banded) – for three different magnitudes of one externally supplied resource (Low Inflow: *ρ*_1_ = 0.001, Medium Inflow: *ρ*_1_ = 0.1 and High Inflow: *ρ*_1_ = 1). The *S* = 15 consumers grow according to our structured Liebig’s law model. All other resources are not supplied (*ρ*_*i*_ = 0 for *i* = 2, …, *S*). In each case, the production network is given by the constant parameterization where all other resources are produced from a given incoming resource. For the banded network, the equilibrium is stable regardless of resource inflow levels, while the sparse network becomes unstable when the resource inflow is low. Parameters: *S* = 15, *C*_*d*_ = 5, *η* = 1, *ϵ* = 0.05 and consumption coefficients sampled from uniform distributions on [0.5, 1.5] before the constraints are imposed.

## Discussion

Unlike many classic ecological models, in which species interactions are completely determined by the abundances of the competing species [32, 33], our model explicitly tracks the dynamics of resources, as have other recent studies of consumer-resource models [7, 34–36]. In varying resource environments, each consumer can have a different effect on another consumer, so consumer and resource abundances together jointly determine species growth rates. Over the course of the dynamics, the per-capita effect of one species on another can change, in contrast to, for example, the generalized Lotka-Volterra model, where these per-capita interactions are fixed in time. In parallel, consumers grow according to Liebig’s law, meaning that each consumer is limited by only a single resource at a time, even though they can deplete other resources. Liebig’s law has been found to accurately describe microbial growth rates, as well as the resource limitation properties of plants, across many different ecosystems [8, 9, 37–39]. It has recently been shown to emerge from a simple model of bacterial metabolism [14]. In this paper, we investigated how the explicit modeling of the consumption and exchange of resources, together with microbial growth governed by Liebig’s law, affected stability in arbitrarily large microbial communities.

Although our model incorporates a more realistic picture of microbial metabolism than previous consumer-resource theory, it is still a highly coarse-grained version of true microbial resource consumption. We interpret the consumed but non-limiting resources as being used to run intracellular processes, but these resources do not contribute to microbial biomass. With this choice, we are modeling a limiting scenario in which the contribution to microbial biomass from the limiting resource is much larger than that of any non-limiting resource, but there may well be microbial growth dynamics for which this is not a good approximation. A simple modification of our model, where some of the consumed but non-limiting resources are lost in internal metabolism, could partially address such a situation. However, allowing non-limiting resources to contribute to consumer biomass greatly complicates our mathematical analysis, and so is beyond the scope of the present work. In our consumption matrix parameterizations, we have also implicitly assumed that each consumer can deplete many of the available resources. Similarly, we have assumed that any resource can be produced from any other. In reality, there are likely stochiometric rules that these matrices must obey [40–42]. For example, recent experiments [19] have shown that resource production networks are approximately, though not precisely, hierarchical. Understanding how the chemical properties of abiotic nutrients and the metabolic strategies of specific bacterial strains constrain the consumption and production networks is therefore an important direction for future work.

Even though our consumer-resource model does not capture the full complexity of microbial metabolism, it still produced stability criteria that refine our understanding of microbial coexistence. We identified a set of symmetric consumption and production networks that guarantee stability as long as intra-specific regulation is large compared to inter-specific interactions. Using simulations, we then showed that interaction networks which violate our condition give rise to unstable, but feasible, fixed points at low consumer abundances. By contrast, recent theory [43, 44] has shown that population abundances do not affect stability in a randomly parameterized Lotka-Volterra model. In the present work, we find the opposite result – given consumption and production networks which do not satisfy our symmetry conditions, some choices of equilibrium consumer abundances generate stable systems, while others do not. At the same time, there are direct parallels between the theory we have developed here, and the theory of pairwise models. In our second stability condition, we derived a relationship between intra-specific regulation and the variability in inter-specific interactions that is reminiscent of May’s classic stability bound [2, 3, 45]. In our tradeoff case, this comparison is particularly direct, with the the mean deviation of the consumption coefficients, which quantifies the variability in the species interactions, playing the role of the interaction heterogeneity in May’s theory [2, 3].

Aside from pairwise models, our theoretical results are also reminiscent of other analyses of model microbial communities. Recent simulations of consumer-resource models with cross-feeding showed that, when consumers are resource limited, the constituent species interact in a characteristic pattern at equilibrium [6]. These results mirror our simulations, where ecosystems with specific symmetric interaction structures are protected from instability at low resource inflow. In [6], the characteristic interaction patterns emerge from community assembly, while in our theory, we impose them from the outset. This connection is particularly interesting because it suggests that special interaction structures can emerge from assembly processes in specific resource environments. Similarly, a recent mathematical analysis of consumer-resource models with multiple forms of consumption also showed that resource inflow mediates a transition to instability [5]. This recent theory, however, treats resource exchange as coming directly from consumer biomass, as though resource production were an additional source of mortality, rather than as an explicit transformation of resources. As a result, the overall strength of production for each species is a tunable parameter and, if it exceeds the total consumption of a single species, then the feasible equilibrium can be unstable [5]. In our model, the strength of production is determined by the resource consumption that is not used for growth, and so cannot exceed the total consumption for each species. Nevertheless, we find that unstable equilibria are possible because of the mismatch between resource depletion and consumer growth. As we have noted before, our theory still applies to the case where we set all of our production coefficients to zero (ie. *P*_*ji*_ = 0), violating the conservation of resource biomass. Then, the system literally has only competitive interactions, but instability is still possible, in direct contrast to the results of [4], where stability is guaranteed because depletion and growth are directly coupled.

Our results can also be seen as a multi-species generalization of classic stability results for species competing for two resources (termed contemporary niche theory) where instability occurs because of the difference between impact and sensitivity vectors [17, 46–48]. In contemporary niche theory, there are three criteria which must be satisfied for a stable equilibrium to exist. First, the species zero net growth isoclines (ZNGIs) must intersect. In our model, this is always true, since each species is limited by a single resource, so it is straightforward to find resource abundances where every consumers’ growth is zero. Second, each species must impact the resources the it finds most limiting more strongly than it impacts other resources. Our second stability criteria is a direct generalization of this result – if each species more strongly regulates the resource they require for growth than they affect all other resources in the system, then the equilibrium is stable. Third, the supply point must lie above the ZNGIs for the coexisting species. In our theory, we ensure that this criteria is satisfied by requiring the equilibrium to be feasible through Eq. (3). We also show in the S1 Appendix that our second stability criterion and the feasibility criteria are closely related – as species more strongly regulate their most limiting resource, the likelihoods of both stability and feasibility are increased. Our theory can be seen as an extension of prior work applying contemporary niche theory to species that interact through the consumption of essential nutrients [17, 49]. This comparison is particularly direct when we remove cross-feeding from our model by setting *P*_*ji*_ = 0, since earlier work did not focus on microbial systems in particular [17]. A similar generalization of this biological scenario found that large numbers of species coexist in oscillatory or chaotic dynamics [50–52], so an important direction for future work will be to better understand the behavior of our model away from equilibrium. It would also be interesting to rigorously understand the stability properties of an ecosystem where consumers grow on many different substitutable resources at variable efficiencies but still leak resources back into the environment through cross-feeding, as in [6]. Last, the model we have considered here is completely deterministic, and so the mismatch between resource depletion and consumer growth resulted from a specific modeling choice. In a stochastic model, depletion and growth may be decoupled through only the differing fluctuations that the consumers and resources undergo, potentially yielding the same stability transition we have observed in our model. More generally, the combination of stochastic drift and the biological mechanisms we have explored here could produce interesting macroecological patterns, as in [53, 54].

By contrast, there is no clear analog of our first stability criterion for low diversity ecosystems. It can, however, be interpreted as perfectly balanced pairwise competition between the consumers, even though the model itself is not built on pairwise competition coefficients. The phenomenon that reciprocity promotes stability has been found in other theoretical studies of microbial communities [5], but also in a diverse set of other fields, from the evolution of cooperation [55,56] to the exchange of food in early societies [29,57]. In addition to our symmetric stability condition, we showed numerically that other consumption and production networks that generate *B* matrices whose eigenvalues are purely real also prevent instability at low consumer abundances. Although we don’t have a precise understanding of how these network structures promote stability, we can describe intuitively how the imaginary parts of the eigenvalues of *B* affect the spectrum of the Jacobian. As the consumer abundances are reduced in our model, the spectrum of the Jacobian begins to have many eigenvalues with non-zero imaginary parts. These complex eigenvalues are separated into two clouds above and below the real axis (see the S1 Appendix for a picture). The imaginary parts of the eigenvalues of *B* control the width of these clouds, while the consumer abundances determine where they are centered. If *B* has purely real eigenvalues, then these two clouds have no width, so regardless of how small the consumer abundances become, no eigenvalues will cross over the imaginary axis. If instead *B* has eigenvalues with non-zero imaginary part, then the clouds in the spectrum of *J* will have non-zero width, and some of these eigenvalues will cross the imaginary axis at a small value of *n*, creating an unstable fixed point. This description also helps to explain why the interaction network *B* need not be precisely symmetric to still promote, but not guarantee, stability. As the imaginary parts of the eigenvalues of *B* become smaller, so too does the width of the eigenvalue clouds in the spectrum of *J*. As a result, networks which are not precisely symmetric but do have eigenvalues with small imaginary parts are better protected from instability at low consumer abundance than those with larger imaginary parts. A more complete mathematical understanding of the connection between the spectrum of *B* and the spectrum of *J* would give us a deeper understanding of why certain modes of resource exchange are stabilizing.

Because our consumer-resource model connects coexistence patterns to empirically accessible quantities, our theoretical results can be tested experimentally. One direct test of our theory would be to design small microbial communities with engineered interaction networks such as those in [58]. Then, an experimentalist can manipulate, for example, the degree to which the interaction network is reciprocal, and observe whether or not coexistence if favored. Conversely, our theoretical results suggest an interpretation of coexistence outcomes when the exact consumption and production networks are unknown. When an experimentalist reduces the resource inflow rates in a serial dilution experiment, the resulting coexistence (or lack thereof) suggests which types of consumption and production networks may be present. Because we showed numerically that our analysis applies to a variety of resource inflow profiles, our theory may also delineate the boundaries of stable coexistence in recent experiments where only one nutrient is externally supplied [18, 19, 59]. For example, recent work has shown experimentally how the resource production network explains the variation in species richness as more resources are externally supplied [19]. Our theory suggests that, if this metabolite production network has eigenvalues with small imaginary parts, the community will be better protected from resource scarcity. Future work should seek to further clarify how the relationship between coexistence outcomes and resource inflow changes depending on network structure. This line of research is especially important because of the difficulty in obtaining well-resolved and quantitative consumer-resource networks for diverse microbial communities.

## S1 Appendix

## Supplementary proofs, calculations and simulation results

We provide proofs of the main results, additional numerical simulations and descriptions of all of the matrix parameterizations.

## Acknowledgments

We thank Seppe Kuehn, Simon Levin and Jonathan Levine for helpful comments and discussion. This material is based upon work supported by the National Science Foundation Graduate Research Fellowship Program under Grant No. DGE-2039656 and Grant No. DGE-1746045. Any opinions, findings, and conclusions or recommendations expressed in this material are those of the author(s) and do not necessarily reflect the views of the National Science Foundation. J.P.O. acknowledges funding from Simons Foundation Grant No. 376199 (www.simonsfoundation.org) and McDonnell Foundation Grant No. 220020439 (www.jsmf.org).

