## Appendix for "Stability criteria for the consumption and exchange of essential resources"

### 1 Stability Criteria

In this section, we derive the sufficient stability criteria that we described in the main text. We show that, if a fixed point exists in the dynamics of our example consumption and production structures and if our stability criteria are satisfied, the fixed point is guaranteed to be stable. We describe how these stability criteria apply to our example consumption and production structures. Last, we discuss under what conditions our sufficient stability criteria are also necessary.

#### 1.1 Deriving the Jacobian

Suppose that the model in Eq. (1) of the main text admits a fixed point with abundances $\vec{R}^\star$ and $\vec{N}^\star$. As discussed in the main text, near to a fixed point, the model reduces to the dynamical system in Equation (2), so we evaluate the Jacobian $\mathbf{J}$ of Eq. (2) at the fixed point $\vec{R}^\star$ and $\vec{N}^\star$ to determine its stability. If the eigenvalues of the Jacobian all have negative real part, then the equilibrium is stable to perturbations. Let's compute the Jacobian of this system.

$$
\begin{aligned}
\frac{\partial \dot{R}_i}{\partial R_j} &= -\delta_{ij} \sum\nolimits_k C_{ik} N_k + P_{ji} \sum\nolimits_k C_{jk}(1-\epsilon_{jk}\delta_{jk}) N_k \\
\frac{\partial \dot{R}_i}{\partial N_j} &= -R_i C_{ij} + \sum\nolimits_k P_{ki} R_k C_{kj}(1-\epsilon_{jk}\delta_{kj}) \\
\frac{\partial \dot{N}_i}{\partial R_j} &= N_k \epsilon_{kk} C_{kk} \\
\frac{\partial \dot{N}_i}{\partial N_j} &= 0
\end{aligned}
\tag{S1}
$$

In matrix notation, the Jacobian is

$$
J = \begin{bmatrix} -[C\vec{N}]_d + P^T[\tilde{C}\vec{N}]_d & -\vec{R}_d C + P^T \vec{R}_d \tilde{C} \\ \vec{N}_d \vec{\epsilon}_d C_{diag} & 0 \end{bmatrix} \tag{S2}
$$

where $C_{diag}$ is the diagonal matrix with entries from the matrix $C$. In the following sections, we will derive the criteria for the stability of the Jacobian $\mathbf{J}$ under some additional assumptions.

#### 1.2 Sufficient conditions for stability

First, we show that $0$ is not an eigenvalue of $J$, so that we can invert the matrix $-\lambda I$ (where $\lambda$ is an eigenvalue of $J$) when applying block determinant rules. Suppose $0$ is an eigenvalue of $J$, then $J$ is singular and there exists a non-zero vector, whose first $S$ components we will denote by $\vec{v}$, and whose second $S$ components we will denote by $\vec{w}$, such that

$$J \begin{bmatrix} \vec{v} \\ \vec{w} \end{bmatrix} = \begin{bmatrix} -[C\vec{N}]_d + P^T[\tilde{C}\vec{N}]_d & -\vec{R}_d C + P^T \vec{R}_d \tilde{C} \\ \vec{N}_d \vec{\epsilon}_d C_{diag} & 0 \end{bmatrix} \begin{bmatrix} \vec{v} \\ \vec{w} \end{bmatrix} = \vec{0}\,. \tag{S3}$$

Since $\vec{N}_d \vec{\epsilon}_d C_{diag}$ is a diagonal matrix with positive entries by assumption, it is an invertible matrix and we know that $\vec{w} = \vec{0}$. Therefore, $\left(-[C\vec{N}]_d + P^T[\tilde{C}\vec{N}]_d\right)\vec{v} = \vec{0}$. In other words, $-[C\vec{N}]_d + P^T[\tilde{C}\vec{N}]_d$ is singular. Because $0 < \epsilon_{ii} \leq 1$, we know that each entry of $[C\vec{N}]_d$ is strictly larger than the corresponding entry of $[\tilde{C}\vec{N}]_d$. As a result, there exists a diagonal matrix $D$ such that $D[C\vec{N}]_d = [\tilde{C}\vec{N}]_d$ and each entry of $D$ is less than 1. We get that $-[C\vec{N}]_d + P^T[\tilde{C}\vec{N}]_d = (-I + P^T D)[C\vec{N}]_d$. $[C\vec{N}]_d$ is invertible since it is a diagonal matrix with non-zero values. Since the column sums of $P^T$ are all $1$ by assumption, the column sums of $P^T D$ are all less than $1$. Therefore, the matrix $-I + P^T D$ is diagonally dominant [1] and hence it is non-singular, implying that $\vec{v} = \vec{0}$, which is a contradiction. We conclude that $0$ is not an eigenvalue of $J$.

Now, we apply block determinant rules to the matrix $J - \lambda I$ and compute the determinant to find that

$$\det[J - \lambda I] = \det[-\lambda I] \det \left[ -[C\vec{N}]_d + P^T[\tilde{C}\vec{N}]_d - \lambda I + \frac{1}{\lambda}[-\vec{R}_d C + P^T \vec{R}_d \tilde{C}]\vec{N}_d \vec{\epsilon}_d C_{diag} \right] \tag{S4}$$

We concentrate on a simplified version of the model, in which $\vec{R^\star} = r\vec{1}$ and $\vec{N^\star} = n\vec{1}$ where $\vec{1}$ is a vector of all $1$'s. We also assume that $C_{ii} = C_d$ and $\epsilon_{ii} = \epsilon$ for all $i$. Let's denote the two relevant matrices in the determinant formula as $A = -[C\vec{N}]_d + P^T[\tilde{C}\vec{N}]_d = -n[C\vec{1}]_d + nP^T[\tilde{C}\vec{1}]_d$ and $B' = [-\vec{R}_d C + P^T \vec{R}_d \tilde{C}]\vec{N}_d \vec{\epsilon}_d C_{diag} = nr\epsilon C_d[-C + P^T \tilde{C}]$. Let's assume that there is an eigenvalue $\lambda$ with Re$(\lambda) > 0$ and look for a contradiction. We will show that, under a specific set of assumptions, $A$ and $\frac{1}{\lambda}B'$ are both negative definite matrices, meaning that their sum is negative definite as well. Since both $A$ and $B'$ are not necessarily symmetric, we actually need to show that the Hermitian parts of $A$ and $\frac{1}{\lambda}B'$ are negative definite. Once we have shown that the Hermitian parts of both $A$ and $\frac{1}{\lambda}B'$ are negative definite, then the Hermitian part of their sum $A + \frac{1}{\lambda}B'$ is also negative definite. Therefore, the assumption that Re$(\lambda) > 0$ is false and we will have found a contradiction.

First, we show that the Hermitian part of $\frac{1}{\lambda}B'$ is negative definite if two conditions are met. We can focus on the simplified matrix $\frac{1}{\lambda}B = \frac{1}{\lambda}[-C + P^T\tilde{C}]$ and ignore the abundances $\vec{R}$ and $\vec{N}$ as well as the diagonal efficiencies $\epsilon_{ii}$ and diagonal consumption coefficients $C_{ii}$ which entered our original formula for $B'$. We assume that the matrix $B$ is symmetric, which is the first stability criterion in the main text. The Hermitian part of $\frac{1}{\lambda}B$ is then:

$$H\left(\frac{1}{\lambda}B\right) = \frac{1}{2||\lambda||^2}\left(\left(\text{Re}(\lambda) - \text{Im}(\lambda)\right)B + \left(\text{Re}(\lambda) + \text{Im}(\lambda)\right)B^T\right) = \frac{\text{Re}(\lambda)}{||\lambda||^2}B\ . \tag{S5}$$

Because $\frac{\text{Re}(\lambda)}{||\lambda||^2}$ is a positive constant by assumption, it does not change the sign of the eigenvalues of $B$. A symmetric matrix is negative definite if and only if all of its eigenvalues are negative, which is the second condition in the main text. Therefore, we have shown that, given the two stability criteria in the main text, the Hermitian part of of $\frac{1}{\lambda}B'$ is negative definite.

Now, we show that the symmetry of $B$ implies that the Hermitian part of $A$ is negative definite. The Hermitian part of $A$ is $H(A) = \frac{1}{2}\left(A + A^T\right) = -[C\vec{N}]_d + \frac{1}{2}\left(P^T[\tilde{C}\vec{N}]_d + [\tilde{C}\vec{N}]_d P\right)$. Let's use the Gershgorin circle theorem to bound the eigenvalues of $H(A)$. The Gershgorin circle theorem states that the eigenvalues of a matrix are contained in discs in the complex plane whose centers are given by the diagonal entries of the matrix with radii given by the sum of the absolute value of the off-diagonal entries of the corresponding row [1]. Therefore, $H(A)_{ii} + \sum_{j\neq i}|H(A)_{ij}| < 0$ for each $i$, the eigenvalues of the Hermitian part of $A$ are all contained in the left half of the complex plane and $H(A)$ is negative definite. The $i$-th row sum of $H(A)$ is

$$\begin{aligned}\sum_j H(A)_{ij} &= -\sum\nolimits_j C_{ij}N_j + \frac{1}{2}\left(\sum\nolimits_j P^T_{ij}\sum\nolimits_k \tilde{C}_{jk}N_k + \left(\sum\nolimits_j \tilde{C}_{ij}N_j\right)\left(\sum\nolimits_j P_{ij}\right)\right) \\ &= \frac{1}{2}\left(-\sum\nolimits_j C_{ij}N_j - \epsilon C_{ii}N_i + \sum\nolimits_j P^T_{ij}\sum\nolimits_k \tilde{C}_{jk}N_k\right) < 0\end{aligned} \tag{S6}$$

where we used the definition of $\tilde{C}$ and the fact that $\sum_j P_{ij} = 1$. Re-organizing and letting $N_i = n$, we find that this Gershgorin inequality is equivalent to $-\sum_j C_{ij} + \sum_k\sum_j P^T_{ij}\tilde{C}_{jk} < \epsilon C_{ii}$. We can then recognize the left-hand side of this inequality as the row sums of the matrix $B$. Since $B$ is symmetric, these row sums are the equal to the row sums of $B^T$. As a result, we have that $B\vec{1} = B^T\vec{1} = -C^T\vec{1} + \tilde{C}^T P\vec{1} = (-C^T + \tilde{C}^T)\vec{1} < 0 < \epsilon C_{ii}$ because the row sums of $P$ are all $1$ from conservation of resource biomass. Therefore, the symmetry of $B$ (the first stability criterion in the main text) implies that the Hermitian part of $A$ is negative definite.

Overall, we have shown that, if our two sufficient stability criteria are satisfied, then the matrices $A$ and $\frac{1}{\lambda}B'$ are both negative definite. The sum of negative definite matrices is also negative definite, so $A + \frac{1}{\lambda}B'$ is negative definite as well. We assumed that $\text{Re}(\lambda) > 0$ but all of the eigenvalues of a negative definite matrix have negative real part, which is a contradiction. Therefore, none of the eigenvalues of $J$ have positive real part if our stability criteria are met. We have not explicitly dealt with the possibility that $\lambda$ is purely imaginary, but our argument shows that this is not possible either. Specifically, the Hermitian part of $B'$ is then zero, and our sufficient stability criteria for $A$ guarantee that the sum $A + \frac{1}{\lambda}B'$ is negative definite once again. We conclude that our two criteria imply that $J$ is stable.

#### 1.3 Gershgorin bound implies that $B$ is negative definite

We now derive the inequality that we stated in the main text which implies that $B$ is stable. It is a straightforward application of the Gershgorin circle theorem to the matrix $B$. If $H(B)_{ii} + \sum_{j \neq i} |H(B)_{ij}| < 0$ for each $i$, the eigenvalues of the Hermitian part of $B$ are all contained in the left half of the complex plane and $H\left(\frac{1}{\lambda}B\right)$ is negative definite. We need that, for each $i$,

$$C_d - \sum_k P_{ik}\tilde{C}_{ik} > \sum_{j \neq i} | - C_{ij} + \sum_k P_{ki}\tilde{C}_{kj}| \tag{S7}$$

to guarantee the negative definiteness of $B$, as we state in the main text. We use this inequality mainly to demonstrate how the diagonal elements of the matrix $B$ can mediate a transition from instability to stability as they become large.

#### 1.4 Stability criteria for the example consumption and production patterns

We now show that the example consumption and production structures which we described in the main text automatically generate symmetric $B$ matrices, satisfying our first sufficient stability criterion. In the tradeoff parameterization, $P_{ij} = \frac{1}{S-1}$ when $i \neq j$ while $P_{ii} = 0$. $C$ is a symmetric matrix with row (or column) sums $\sum_j C_{ij} = C_d + (S-1)C_0$ for each $i$. The $(i,j)$-th entry of the matrix product $P^T\tilde{C}$ is $[P^T\tilde{C}]_{ij} = \frac{1}{S-1}\left(C_d + (S-1)C_0 - C_{ij}\right)$, while the $(i,i)$-th entry is $[P^T\tilde{C}]_{ji} = \frac{1}{S-1}\left(C_d + (S-1)C_0 - C_{ji}\right)$. Since $C$ is symmetric, $C_{ij} = C_{ji}$ and $P^T\tilde{C}$ is symmetric as well. Therefore, $B$ is symmetric since it is the sum of symmetric matrices, and the first stability condition is satisfied. Now, we compute both sides of the Gershgorin inequality that implies the second stability criterion in this parameterization. We get that $C_d - \sum_k P_{ik}\tilde{C}_{ik} = C_d - C_0$ and that

$$\begin{aligned} \sum_{j \neq i} | - C_{ij} + \sum_k P_{ki}\tilde{C}_{kj}| &= \sum_{j \neq i} \left| -C_{ij} + \frac{1}{S-1}\left((1-\epsilon)C_d + (S-1)C_0 - C_{ij}\right)\right| \\ &\geq \sum_{j \neq i} |C_0 - C_{ij}| + \frac{1}{S-1}\sum_{j \neq i} |(1-\epsilon)C_d - C_{ij}| \\ &\geq \sum_{j \neq i} |C_0 - C_{ij}| + (1-\epsilon)C_d + C_0 \end{aligned} \tag{S8}$$

where we used the triangle inequality twice. Combining these, we get that if $\epsilon C_d > 2C_0 + \sum_{j \neq i} |C_0 - C_{ij}|$, then the second stability criteria is true, as in Equation (4) in the main text. Note that this bound is not as tight as the inequality derived using the Gershgorin circle theorem or the second stability criterion itself.

Now, we consider the symmetric circulant matrix case. Symmetric circulant matrices have constant row and column sums. For consistency with the constant sums case, we let $\sum_j C_{ij} = C_d+(S-1)C_0$ for each $i$. For the first stability criterion, circulant matrices commute with one another, so if $C$ and $P^T$ are both symmetric and circulant, we have that $(P^TC)^T = C^TP = CP^T = P^TC$. In other words $P^TC$ is symmetric and therefore so is $B = -C + P^TC$ and the first condition is always satisfied. It is possible for the second condition to be false, so we need to explicitly check this condition, as we do in the main text.

##### 1.5 Stability criteria without cross-feeding

When there is no cross-feeding, $P_{ij} = 0$ for all of its entries. In this section, we assume that $\sum_{j\neq i} C_{ij} = (S-1)C_0$ for analytical tractability, and we also continue to assume that $\vec{R}^\star = r\vec{1}$, $\vec{N}^\star = n\vec{1}$, $C_{ii} = C_d$ and $\epsilon_{ii} = \epsilon$ for all $i$. The eigenvalues $\lambda$ of the Jacobian $J$ satisfy the characteristic polynomial

$$0 = \det(J-\lambda I) = \det\left(\begin{bmatrix} -[C\vec{N}]_d - \lambda I & -\vec{R}_dC \\ \vec{N}_d\epsilon_dC_{diag} & -\lambda I\end{bmatrix}\right) = \det\left(\begin{bmatrix} -n((S-1)C_0+C_d)I - \lambda I & -rC \\ nC_dI & -\lambda I\end{bmatrix}\right). \tag{S9}$$

First, let's decide if $\lambda = -n((S-1)C_0+C_d)$ can be an eigenvalue. Let $v, w \in \mathbb{R}^S$ and suppose that

$$\begin{bmatrix} 0 & -rC \\ n\epsilon C_dI & n((S-1)C_0+C_d)I\end{bmatrix}\begin{bmatrix} v \\ w\end{bmatrix} = \vec{0}\,. \tag{S10}$$

Since $-rC$ is invertible, $w = \vec{0}$, but then $v = \vec{0}$ as well, and $\lambda = -n((S-1)C_0+C_d)$ is not an eigenvalue of $J$. Therefore, we can safely apply block determinant rules to find all the eigenvalues. We get that

$$0 = \det\left(-\vec{N}_d\epsilon_dC_{diag}\left([C\vec{N}]_d+\lambda I\right)^{-1}\vec{R}_dC - \lambda I\right) = \det\left(-\frac{nr\epsilon C_d}{n((S-1)C_0+C_d)+\lambda}C - \lambda I\right) \tag{S11}$$

so we can write down the eigenvalues of $J$ in terms of the eigenvalues of $C$. If $\omega$ is an eigenvalue of $C$, then the roots of the polynomial

$$\lambda^2 + n((S-1)C_0+C_d)\lambda + nr\epsilon C_d\omega = 0 \tag{S12}$$

are eigenvalues of $J$. So, we want to know when the roots of this complex quadratic have negative real part. We can derive constraints on $\omega$ under which $\lambda$ is guaranteed to have negative real part by using the quadratic formula and the identity $Re\left(\sqrt{x+iy}\right) = \frac{1}{\sqrt{2}}\sqrt{\sqrt{x^2+y^2}+x}$, or just by appealing to the Routh-Hurwitz stability criteria. In either case, if

$$\begin{aligned} &n((S-1)C_0 + C_d) > 0 \\ &n((S-1)C_0 + C_d)^2 Re(\omega) > rd Im(\omega)^2 \end{aligned} \tag{S13}$$

hold, then $Re(\lambda) < 0$. The first constraint in (S13) is always satisfied from the definitions. The second constraint is the interesting one. First of all, if $Re(\omega) \leq 0$ (ie. if $-C$ is unstable), then the Jacobian $J$ is unstable, since the right hand side of (S13) is non-negative. So, we can already rule out consumption structures ($C$ matrices) that are unstable by themselves. In addition to the stability of $C$, (S13) shows that the magnitude of $Im(\omega)$ must be small compared the magnitude of $Re(\omega)$. If $Im(\omega) \neq 0$, which is true for a generic consumption matrix, then by sending $n \to 0$, we will eventually get instability. These results directly mirror those presented in the main text, where low consumer abundances lead to instability. Moreover, the criteria we have derived in this simpler case unify our two stability criteria in the more general case. To see this, let's interpret our more general stability criteria in the non-cross-feeding parameterization. The first symmetry criterion in the general case forces the imaginary part of the eigenvalues of $C$ to be zero ($Im(\omega) = 0$) and the second stability criterion ensures that the matrix $C$ is itself stable. In the non-cross-feeding case, however, we see that these stability criteria are restrictive. It is possible to have stability without necessarily being precisely symmetric (and therefore forcing $Im(\omega) = 0$), as long as the real parts of the eigenvalues of the matrix $C$ are large enough. These results mirror our conjecture in the main text that if the matrix $B$ has small imaginary parts, then it will be protected from stability at low consumer abundances (see also Fig. H).

#### 1.6 Calculating the eigenvalues of $J$ when $A$ and $B$ are symmetric and simultaneously diagonalizable

In the previous section, we showed how the eigenvalues of $J$ are directly related to the eigenvalues of $C$ when there is no cross-feeding. Now, we prove a similar result in a different context. We show that the eigenvalues of $J$ can be written in terms of the eigenvalues of the matrices $A$ and $B$ provided that these matrices are simultaneously diagonalizable. In particular, the two consumption and production structures which we present in the main text (the tradeoff and circulant cases) give rise to $A$ and $B$ matrices which are simultaneously diagonalizable.

In the proof of our sufficient stability criteria, we showed that each non-zero eigenvalue $\lambda$ of $J$ satisfies

$$0 = \det\left[\lambda A + n\epsilon r C_d B - \lambda^2 I\right] \tag{S14}$$

which means that the matrix $\lambda A + n\epsilon r C_d B - \lambda^2 I$ has an eigenvalue of $0$. Equivalently, there exists an vector $\vec{v}$ such that

$$\lambda A\vec{v} + n\epsilon r C_d B\vec{v} = \lambda^2\vec{v} \tag{S15}$$

for each eigenvalue $\lambda$ of $J$. When $A$ and $B$ are simultaneously diagonalizable, they share the same eigenspaces, so we can find a vector $\vec{v}$ that is an eigenvector of both $A$ and $B$. Let $\lambda_A$ denote the corresponding eigenvalue of $A$ and $\lambda_B$ the corresponding eigenvalue of $B$. Then, we can re-write the above vector equation in terms of these eigenvalues and solve for $\lambda$ to get that

$$\lambda = \frac{1}{2}\left(\lambda_A \pm \sqrt{\lambda_A^2 + 4n\epsilon r C_d \lambda_B}\right) \tag{S16}$$

so we have solved for the eigenvalues of $J$ in terms of the corresponding pairs of eigenvalues of $A$ and $B$. Using Eq. S16, we can also prove that $J$ is stable if and only if $B$ is stable when $B$ is symmetric. If $A$ and $B$ are symmetric, then both $\lambda_A$ are $\lambda_B$ are real. In this case, $\sqrt{\lambda_A^2 + 4n\epsilon r C_d \lambda_B} \leq \lambda_A$ as long as $\lambda_B < 0$, so that $\lambda < 0$ and the equilibrium is stable. Since $B$ is symmetric, we know that $\lambda_A < 0$, so the only way to have a positive $\lambda$ is to have $\lambda_B > 0$. In other words, the sign of $\lambda$ is simply given by the sign of $\lambda_B$ and $J$ is stable if and only if $B$ is stable. In Fig. I, we plot the predictions of Eq. S16 as well as the spectrum of $J$ and the agreement is exact.

##### 1.7 Is $J$ stable if and only if $B$ is stable when $B$ is symmetric?

We have shown in two different simplified scenarios that $J$ is stable if and only if $B$ is stable provided that $B$ is symmetric, which suggests that this may be true in general. We cannot prove this statement in its full generality. In this section, we first provide additional numerical evidence that it is true. We consider a more general parameterization of the Jacobian:

$$J = \begin{bmatrix} A & B \\ nI & 0 \end{bmatrix} \tag{S17}$$

where $A$ is an arbitrary negative definite matrix, $B$ is an arbitrary symmetric matrix and $n$ is some positive constant. Note that these new definitions of $A$, $B$ and $n$ slightly abuse the notation we have developed because they absorb additional constants (such as $\epsilon$ and $r$), but this choice does not affect any of the qualitative results. This more general parameterization of $J$ includes the Jacobian we have analyzed, but it removes the relationship between $A$ and $B$ that exists in our model. Instead, $A$ and $B$ are arbitrary matrices, as long as they are negative definite and symmetric respectively. In Fig. J, we plot the maximum eigenvalue of $B$ against the real part of the eigenvalue of $J$ with largest real part for $A$ and $B$ matrices with randomly sampled entries and for a few different system sizes $S$. We vary the diagonal entries of $B$ to create stable and unstable matrices (see the R code on Github at https://github.com/tgibbs-hub/essential-stability-criteria for details). Across many replicates, the sign of the leading eigenvalue of $B$ predicts exactly the sign of the leading eigenvalue of $J$ even in this more general parameterization of the Jacobian.

We have already proven that, if $B$ is stable, then $J$ is stable when $B$ is symmetric. One way to prove the reverse implication is to show that, if $B$ is unstable, then so is $J$. We will actually show that, if $B$ is unstable with an odd number of positive eigenvalues and no repeated eigenvalues, then $J$ is unstable. Since $B$ has no repeated eigenvalues, it admits an eigendecomposition. Therefore, there exists a matrix $Q$ such that $B = Q^{-1}\Lambda Q$ where $\Lambda$ is a diagonal matrix containing the eigenvalues of $B$. Because $B$ is symmetric, we also know that $Q$ is orthogonal so that $Q^{-1} = Q^T$. Now, let's set

$$U = \begin{bmatrix} Q & 0 \\ 0 & Q \end{bmatrix} \tag{S18}$$

so that we can define the matrix

$$J_1 = U^{-1}JU = \begin{bmatrix} Q^{-1}AQ & \Lambda \\ nI & 0 \end{bmatrix} . \tag{S19}$$

$J_1$ is similar to $J$, so they share the same eigenvalues. Let's denote the eigenvalues of $B$ by $\lambda_1 > \lambda_2 > \cdots > \lambda_k > 0 > \lambda_{k+1} > \cdots > \lambda_S$ for some odd integer $k \in \{1, \ldots, S\}$ and let's define the matrix $|\Lambda|$ as the entry-wise absolute value of the matrix $\Lambda$. Let's also define the matrix $T$ as a $2S \times 2S$ block matrix with $S \times S$ size blocks. The off-diagonal blocks of $T$ are all zeroes, the upper left-hand block is the identity and the lower right-hand block is a diagonal matrix with $-1$ in the first $k$ columns and $1$ in the remaining columns. Then, we can write the matrix $J_1$ as

$$J_1 = \begin{bmatrix} Q^{-1}AQ & -|\Lambda| \\ nI & 0 \end{bmatrix} T = J_2 T \tag{S20}$$

where we have defined $J_2$. Last, let's define

$$J_3 = UJ_2U^{-1} = \begin{bmatrix} A & Q\left(-|\Lambda|\right)Q^{-1} \\ nI & 0 \end{bmatrix} . \tag{S21}$$

We have already shown that $J_3$ is stable because it satisfies our two stability criteria. In particular, $Q|\Lambda|Q^{-1}$ is symmetric because $Q$ is orthogonal: $[Q\left(-|\Lambda|\right)Q^{-1}]^T = Q^{-T}\left(-|\Lambda|\right)Q^T = Q\left(-|\Lambda|\right)Q^{-1}$. It is also similar to the matrix $-|\Lambda|$ so it has negative eigenvalues and is negative definite. Similarly, $A$ is negative definite by assumption. Therefore, $J_3$ is stable and so is $J_2$ because they are similar. On the other hand, $\det[J_2] = -\det[J_1]$ because $\det[T] = -1$ since $k$ is odd. From this, we can conclude that $J$ has at least one positive real eigenvalue. Since $J$ is a real matrix, its complex eigenvalues come in complex conjugate pairs. The determinant is the product of the eigenvalues, so these conjugate pairs do not change its sign, regardless of whether or not they have positive or negative real part, since they enter the determinant only through their modulus. Since $J_1$ and $J_2$ have opposite signs, we know that at least one of the real eigenvalues of $J_1$ must be positive. If this were not true, then $J_1$ and $J_2$ would have determinants with the same sign. So, because $J$ is similar to $J_1$, $J$ is unstable as well.

Although we have not shown the if and only statement in full generality, we have shown that the transition from stability to instability is sharp when $B$ does not have degenerate eigenvalues. Let's consider slowly changing some parameter (like $C_d$) such that $B$ just becomes unstable. Then, $B$ has only one positive eigenvalue because it does not have any degenerate eigenvalues. From the above argument, we know that $J$ is also unstable. We cannot analytically rule out the possibility that $J$ becomes stable after more of the eigenvalues of $B$ become positive, though we do not observe this behavior in simulations.

### 2 Feasibility Analysis

Throughout the stability analysis in the previous section, we assumed that a fixed point $\vec{R}^{\star}$ and $\vec{N}^{\star}$ existed. We now evaluate when such a fixed point exists, and describe how our feasibility criteria relate to our stability criteria.

#### 2.1 General feasibility criteria

As we stated in the main text, there is a fixed point in the simplified model (where consumers do not change their resource preferences as a function of resource availability) when the abundances

$$\vec{N} = \left[\vec{R}_d C - P^T \vec{R}_d \tilde{C}\right]^{-1} \vec{\rho} \tag{S22}$$

are all positive for the specified resource inflow $\vec{\rho}$. We can solve for the resource abundances immediately because the consumer dynamics involve only one resource. We get that $R_i = \frac{\eta_i}{\epsilon_{ii} C_{ii}}$.

#### 2.2 Feasibility for the example consumption and production patterns

We now show that the abundances $\vec{N} = n\vec{1}$ is a fixed point for the example consumption and production structures that we consider in the main text. We need the vector

$$\vec{\rho} = nr\left[C - P^T \tilde{C}\right]\vec{1} \tag{S23}$$

to have all positive components. In other words, we need the row sums of the matrix $-B = C - P^T\tilde{C}$ to have all positive entries. However, in each of our parameterizations, both the $C$ and $P$ matrices have constant row and column sums. Therefore, $P^T\tilde{C}$ also has equal row sums, and so does $-B$. Therefore, the constant abundances case $n$ gives rise to a constant supply vector $\vec{\rho} = \rho\vec{1}$ where $\rho = C_d + (S-1)C_0 - (1-\epsilon)C_d - (S-1)C_0 = \epsilon C_d$. This constant supply vector is always positive by definition, so the example consumption and production structures are always feasible in our stability analysis. For more complex resource inflows, however, $\vec{N}$ is not a constant vector and we have not proven that these interaction structures guarantee feasibility. Moreover, the randomly sampled consumption and production structures can give rise to feasible or unfeasible abundances, even for the constant abundances case (see Fig. E).

#### 2.3 Sufficient stability criteria imply feasibility

The Gershgorin bound that implies the second stability criterion is $C_d - \sum_k P_{ik}\tilde{C}_{ik} > \sum_{j\neq i} |-C_{ij} + \sum_k P_{ki}\tilde{C}_{kj}|$ for all $i$ which is the same as having the row sums of the matrix $\bar{B}$ be negative, where $\bar{B}$ is the same as the matrix $B$ except with the absolute value of the off-diagonal entries rather than their original values. For feasibility in the case of constant abundances, we need the row sums of the matrix $-B$ to be positive. Therefore, the second stability criterion is a more restrictive condition than feasibility in this case. In Fig. E, we can see the affect of feasibility on the different matrix parameterizations as we vary the parameter $\epsilon$. In the tradeoff and circulant cases, we proved in the previous section that these matrices are always feasible. As a result, the points which are the numerically determined stability thresholds do not change as $\epsilon$ changes, while the Gershgorin bound (the dashed lines in the figure) does change, because it is sensitive to changes in $\epsilon$. For the random case, both the numerically computed thresholds and the Gershgorin bounds change as a function of $\epsilon$. In fact, the analytical bound seems to predict the average numerically determined value fairly well. This is because we enforce both feasibility and stability when we compute the numerical thresholds in Fig. E, so the analytical bound is capturing the behavior of the probability of feasibility at low $\epsilon$, rather than the probability of stability. By contrast, in Fig. D, we only enforce stability in the numerically determined $C_d$ values, so the difference between Fig. D and Fig. E reveals the influence of feasibility on the system.

#### 2.4 Feasibility for different growth rules

Thus far, we have only discussed feasibility in the context of the simplified model where consumers do not change their preferred resource. However, given a feasible fixed point in this simpler model, it is not necessarily achieved in the dynamics of the model with a specified growth rule. For example, there could be a feasible fixed point in which a given consumer actually derives less benefit from a different resource than the one it is consuming. Under Liebig's law, the consumer switches to this resource, meaning that the feasible fixed point for the simplified model is not an equilibrium of the Liebig model. Taking the unstructured version of Liebig's law as an example, we need

$$\min_{j\in\{1,\ldots,S\}}\{\epsilon_{jk}C_{jk}R_j : C_{jk}\neq 0\} = \epsilon_{ii}C_d \tag{S24}$$

for the assignment of consumer to resources to generate the feasible fixed point. In this equation, the $\epsilon$ matrix, in combination with the consumption coefficients and the resource abundances, determines which resources are limiting for each species. Therefore, if the the diagonal efficiencies $\epsilon_{ii}$ are small relative to the off-diagonal efficiencies, then the diagonal resources are limiting for the Liebig law growth rule. In Fig. 5 of the main text and in Fig. F-G, we fix $\epsilon$ and vary $C_d$ which means that the minimum above is eventually realized by a different assignment of consumers to resources. For the structured version of Liebig's law, the upper limit of feasible $C_d$ values is larger since the minimum is over fewer entries in the consumption matrix. For a growth rule with a maximum instead of a minimum, the dependence is reversed. In this case, because $C_d$ is large compared with the remaining consumption coefficients in order to ensure stability, the maximum is automatically realized by the diagonal consumer-resource pairs in the growth dynamics, and the $\epsilon$ matrix does not need to have a small diagonal.

### 3 Matrix Parameterizations

In this section, we summarize the different matrix parameterizations we used to generate Fig. F-G which test our hypothesis that the imaginary parts of the eigenvalues of $B$ control stability. All the code used to run simulations and generate the figures is available on GitHub at https://github.com/tgibbs-hub/essential-stability-criteria. In each case, we set the diagonal values of the consumption coefficients to be $C_d$. We also set the diagonal values of the $P$ matrix to be zero, except for in the upper triangular case, which we describe in more detail below. In Fig. B, we visualize all of the consumption and production matrices we did not display in the main text.

#### 3.1 Consumption matrices

Tradeoff: $C$ is a symmetric matrix with row (or column) sums given by $\sum_j C_{ij} = C_d + (S-1)C_0$ for each $i$. To produce this parameterization numerically, we generate a random symmetric matrix from the underlying uniform distribution. Then, we find the closest matrix with constant row sums using the R package Spbsampling [2].

Circulant: First, we sample a random vector of length $S$ from the underlying uniform distribution. Then, we assign the matrix elements in the permuted fashion described in the main text. Last, we make the matrix symmetric by taking its Hermitian part.

Random: We randomly sample all the off-diagonal entries from the underlying uniform distribution.

Banded: We randomly sample the off-diagonal entries displaced from the diagonal by one index from the underlying uniform distribution. Then, we constrain each row to have a row sum of $C_0$. Therefore, the matrix $B$ when $P$ is constant has only real eigenvalues but is not symmetric.

Correlated: We specify a parameter $p$ which controls the degree of correlation between each off-diagonal pair $(C_{ij}, C_{ji})$. For each pair, we sample one random value from the specified underlying uniform distribution, which we will call $M_{ij}$. Then, $C_{ij} = pM_{ij}+(1-p)\bar{C_{ij}}$ and $C_{ji} = pM_{ij}+(1-p)\bar{C_{ji}}$ where $\bar{C_{ij}}$ and $\bar{C_{ji}}$ are new samples from the underlying uniform distribution. When $p = 1$, $C$ is the symmetric matrix given by $A$, while for $p = 0$, it is fully random.

Lower Triangular: We sample all the off-diagonal elements of $C$ fully randomly from the underlying uniform distribution, and then we set the upper triangular part of the matrix to zero.

Non-symmetric Tradeoff: We sample all the off-diagonal elements of $C$ fully randomly from the underlying uniform distribution, and then we constrain all the row sums to be given by $(S-1)C_0 + C_d$.

Sparse: We randomly select $\lfloor S/3 \rfloor$ off-diagonal entries from each row to be non-zero. Then, we sample these non-zero entries from the underlying uniform distribution.

##### 3.2 Production matrices

Constant: As described in the main text, we set $P_{ij} = 1/(S-1)$ when $i \neq j$ and set $P_i i = 0$

Circulant: We randomly sample a vector of length $S$ from a uniform distribution on $[0, 1]$ and create a circulant matrix by permuting this vector one index as we descend the rows. Then, we set $P_i i = 0$ and make the matrix symmetric by taking the Hermitian part. Last, we multiply the matrix by a constant to make all the row sums equal to $1$.

Random: We sample all the off-diagonal entries from the uniform distribution on $[0, 1]$ and then enforce that all row sums are $1$.

Upper Triangular: We randomly sample all the upper triangular entries from a uniform distribution on $[0, 1]$ and then constrain the row sums to be $1$. This is the only $P$ matrix with non-zero diagonal entries, because otherwise the last resource would not be transformed into anything and conservation of biomass would be violated.

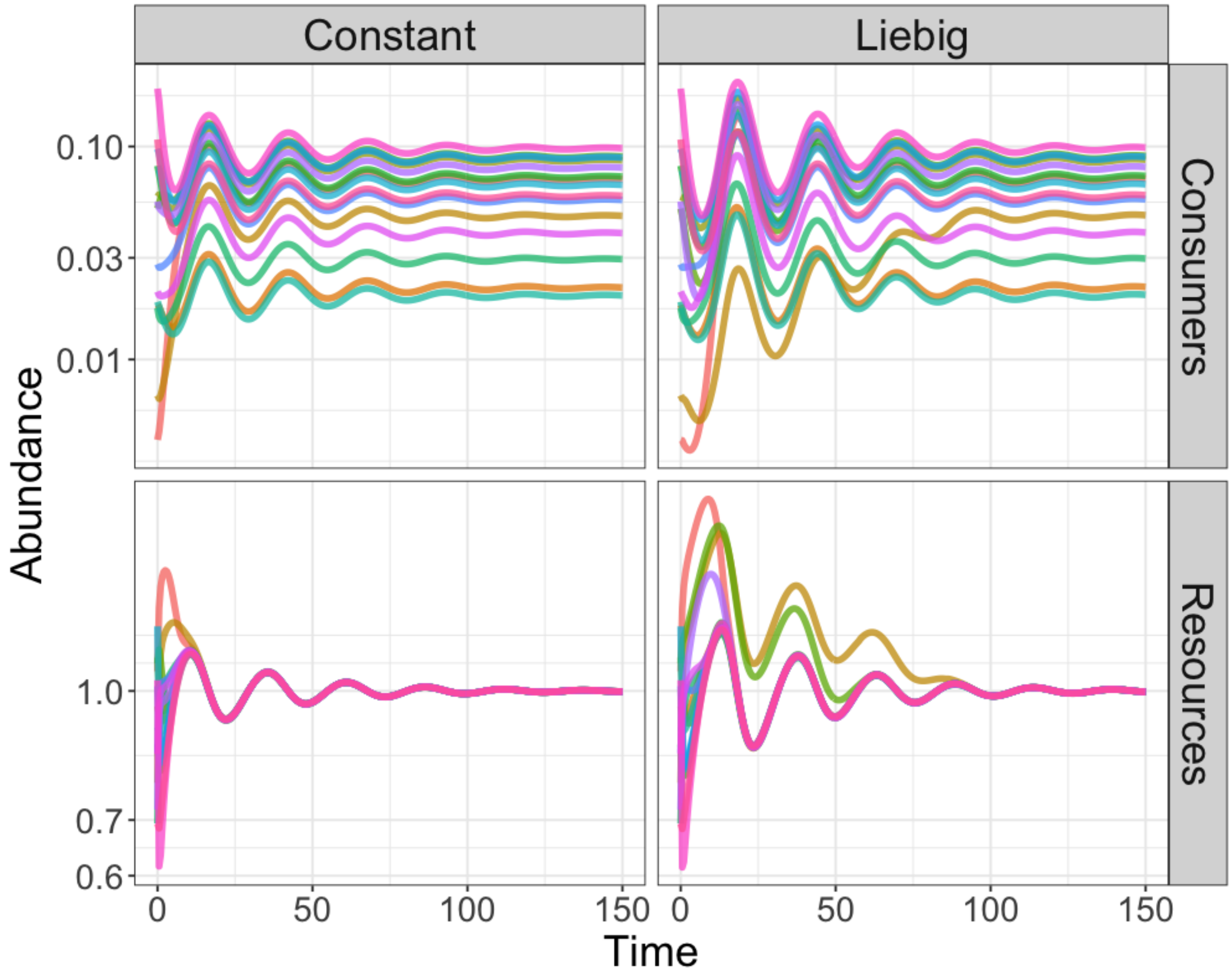

Figure A: **Simplified and Liebig growth rules produce the same equilibria.** We plot the dynamics of consumers and resources (the rows) for two different growth rules (the columns). In the column labeled constant, the consumers never deviate from their assigned resource, while in the column labeled Liebig, they grow according to the unstructured Liebig law growth rule described in the main text. After a brief transient where the consumer and resource abundances differ, both of these models reach the same equilibrium abundances. Parameters: $S = 15$, $C_d = 20$, $\epsilon = 0.05$ and $\vec{\eta} = \vec{1}$. $\rho$ is randomly sampled from a uniform distribution on $[0, 0.1]$. The off-diagonal elements of $C$ are sampled from a uniform distribution on $[1, 3]$ and the off-diagonal elements of $P$ are sampled uniformly and then the row sums are constrained to be $1$.

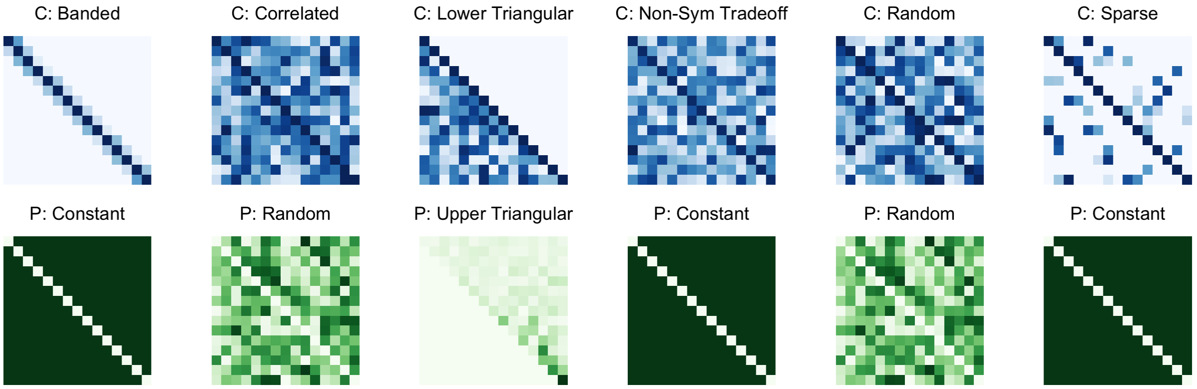

Figure B: **Visualizations of the different matrix parameterizations.** Darker colors indicate larger values. Matrices in the same column are used together to parameterize the system in Fig. F-G, even though we label the heatmaps only by their consumption matrix in Fig. F-G.

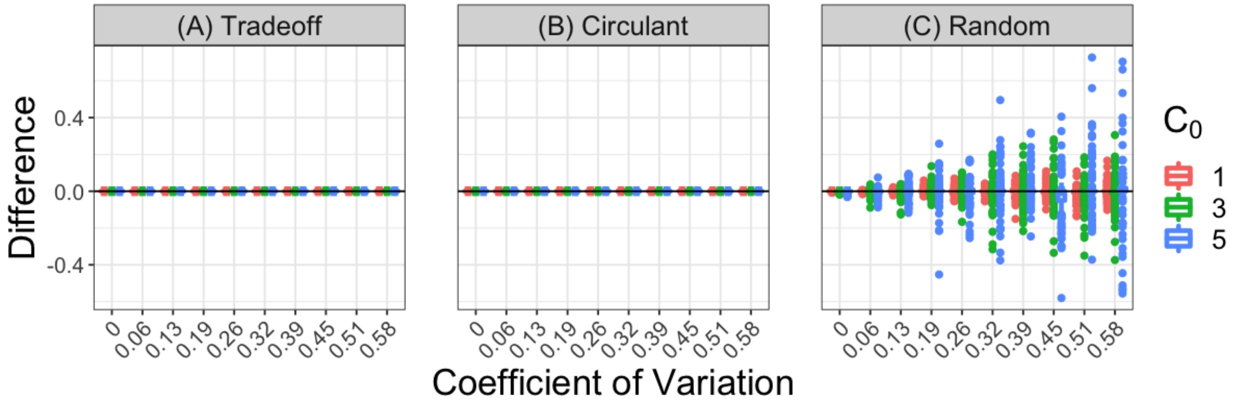

Figure C: **Differences between analytical and numerical $C_d$ values which induce stability.** We plot the difference between the analytical prediction for $C_d$ from the second stability criterion and the smallest value of $C_d$ at which the system becomes stable numerically. We plot the differences as boxplots across a range of different coefficients of variation and two different $C_0$ values (blue and red colors) for the different parameterizations of the consumption and production matrices ((A) is the tradeoff parameterization, (B) is the circulant parameterization and (C) is the random parameterization). The differences are precisely zero when our first criterion is satisfied (panels (A-B)), but in the random case, the difference can be positive or negative. Parameters: $S = 15$, $n = r = 1$ and $\epsilon = 0.5$ for all panels. For each of the different matrix parameterizations, we first sample the consumption coefficients from uniform distributions with mean $C_0$ and the specified standard deviations so that the coefficients of variation vary from $0.001/\sqrt{3}$ to $1/\sqrt{3}$. Then, we impose the constraints for the tradeoff and circulant parameterizations afterwards.

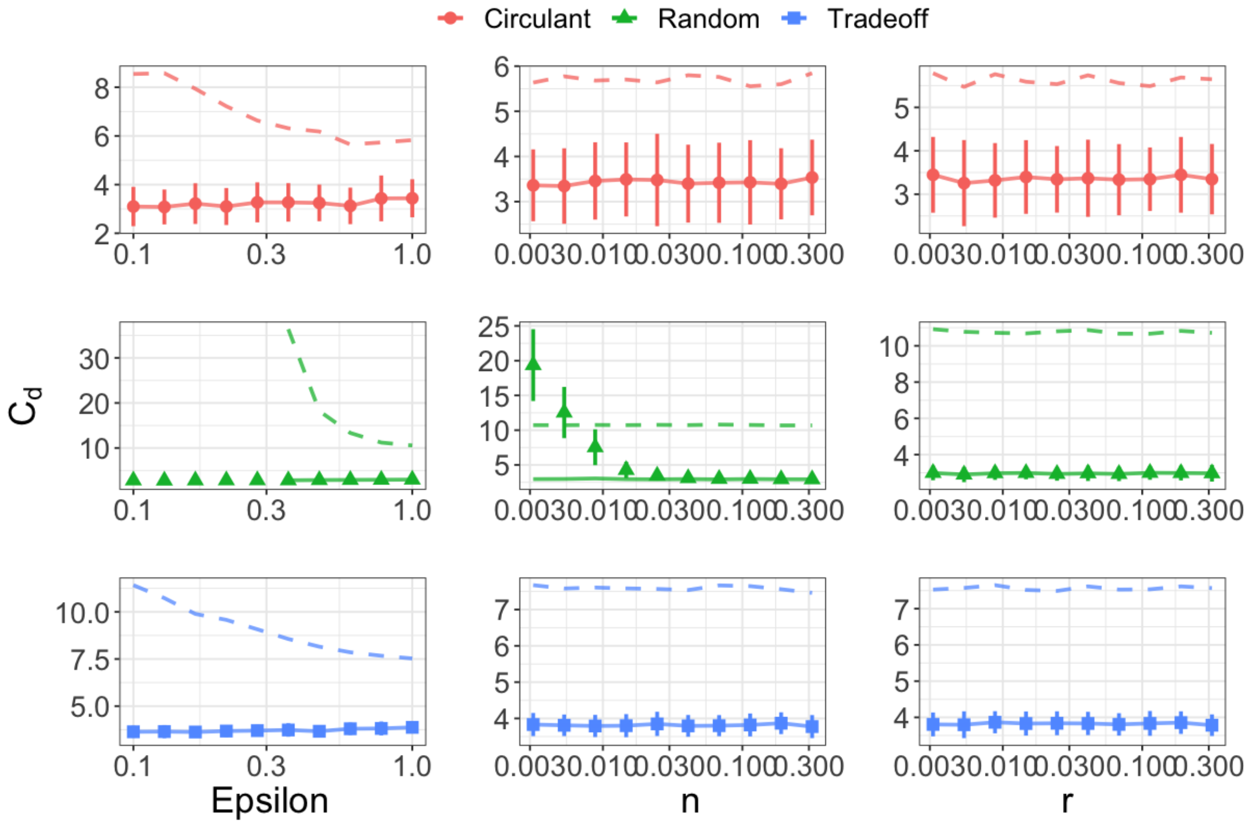

Figure D: **Diagonal consumption $C_d$ required for stability as a function of the parameters $\epsilon$, $n$ and $r$.** We plot the behavior of the smallest $C_d$ at which $B$ becomes stable (the solid lines) and of the Gershgorin bound on $C_d$ (the dashed lines) as well as the numerically determined $C_d$ at which the system first becomes stable (points and error bars) as a function of the different parameters in the model. We consider the three example consumption and production structures. The rows are the different interaction structures and the columns are the different varying parameters. The middle column (where $n$ is changing) is the same as in the main text. We see that the Gershgorin bound is an overestimate of the numerical $C_d$ values, except for small $n$ in the random case, where it is also violated by the empirical values. In the first column, changing $\epsilon$ affects the Gershgorin bound in all three cases, but it does not appreciably change the analytical prediction or the empirical $C_d$ values. This is because we are only enforcing stability in these plots which is not affected by $\epsilon$. In the third column, varying $r$ has no affect on the numerically computed thresholds or the analytical bounds for any of the three matrix parameterizations. Parameters: $S = 15$, the off-diagonal coefficients of $C$ are sampled from a uniform distribution on $[0, 2]$ before each of the matrix parameterizations are imposed. For the $\epsilon$ column, $n = r = 1$. For the $n$ column, $r = 1$ and $\epsilon = 0.9$. For the $r$ column, $n = 1$ and $\epsilon = 0.9$.

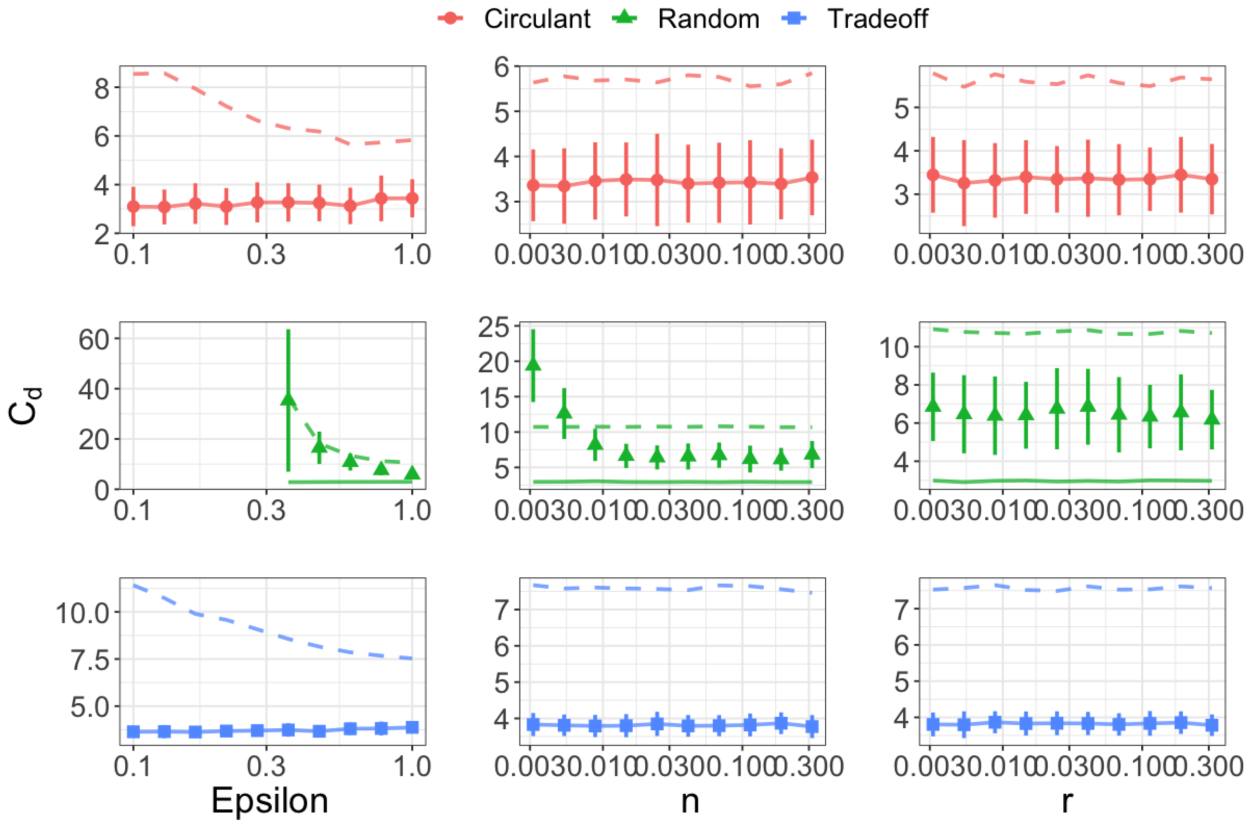

Figure E: **Diagonal consumption $C_d$ required for feasibility and stability as a function of the parameters $\epsilon$, $n$ and $r$.** We plot the behavior of the smallest $C_d$ at which $B$ stable (the solid lines) and of the Gershgorin bound on $C_d$ (the dashed lines) as well as the numerically determined $C_d$ at which the system first becomes feasible *and* stable (points and error bars) as a function of the different parameters in the model. We consider the three example consumption and production structures. The rows are the different interaction structures and the columns are the different varying parameters. The middle column (where $n$ is changing) is the same as in the main text. The feasibility constraint makes the empirical values of $C_d$ larger than those predicted by the second stability criterion because the system becomes stable before it becomes feasible. The Gershgorin bound, however, is still larger than the empirical values (except for small $n$ values) because it is related to the feasibility criterion. In the first column, changing $\epsilon$ affects the Gershgorin bound in all three cases, but it does not appreciably change the empirical values for the circulant or tradeoff cases. In contrast, $\epsilon$ does affect the value of $C_d$ in the random case. These behaviors can once again be explained by the relationship between the sufficient stability criterion and the feasibility criteria, since small values of $\epsilon$ make feasibility less probable. In the third column, varying $r$ has no affect on the numerically computed thresholds or either of the analytical bounds for any of the three matrix parameterizations. Parameters are the same as in Fig. D

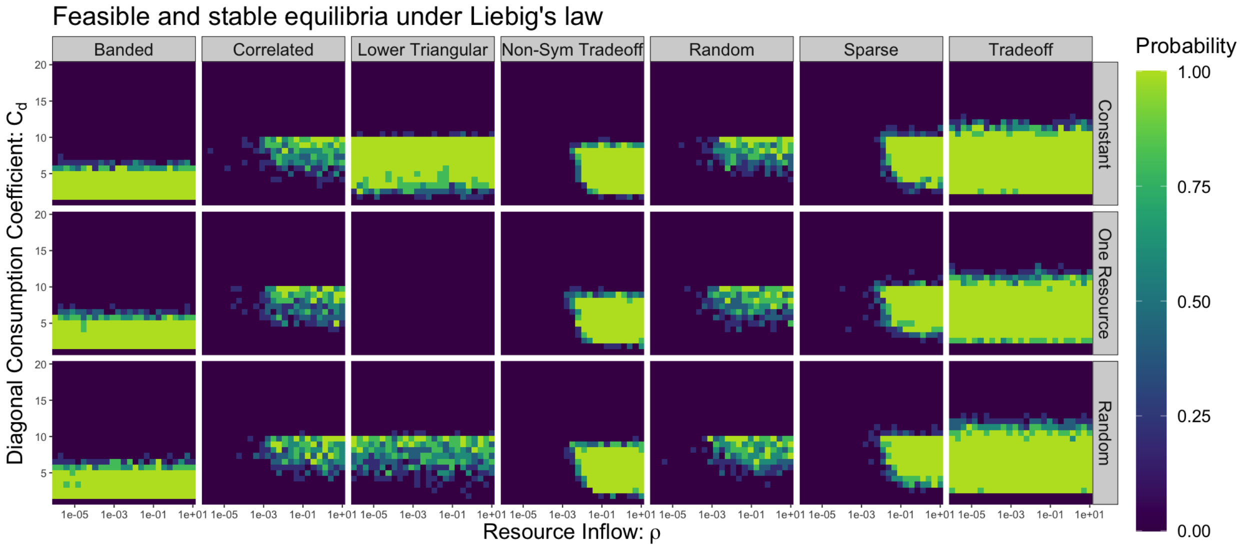

Figure F: **Probability of feasible and stable equilibria under Liebig's law.** We plot the probability of finding a feasible and stable fixed point in $5$ replicates across a range of $C_d$ values and resource inflows $\rho$. We use three different possible resource inflow profiles (the rows). The constant inflow has all resources supplied equally, the one resource inflow supplies only the first resource and the random inflow has all resources supplied at rates sampled from a uniform distribution. In all cases, we ensure that the total resource supply to be given by $\rho$. We enforce that the fixed point is realized under the Liebig's law dynamics, where each consumer grows on the most limiting nutrient of all the resources. The columns are the different consumption matrix parameterizations, as shown in Fig. B. Fig. B also shows the corresponding production matrix used to generate the heatmap. As we described in the main text, the matrix parameterizations which generate $B$ matrices with eigenvalues that have non-zero imaginary parts show a transition to instability at low resource inflow. In contrast, parameterizations whose $B$ matrices have purely real eigenvalues do not show this transition. In the non-symmetric tradeoff and sparse cases, there is a more clear relationship between the transition to instability and the value of $\rho$ when compared to the fully random cases. In particular, systems with larger diagonal consumption $C_d$ tend to be stable at lower levels of resource inflow $\rho$. In each of these cases, the resource inflow profile does not significantly change the probability of feasible and stable fixed points, except for the lower triangular case, which can never be feasible for one externally supplied resource. Parameters: $S = 15$, $\epsilon = 0.05$ and consumption coefficients sampled from uniform distributions on $[0.5, 1.5]$ before the constraints are imposed.

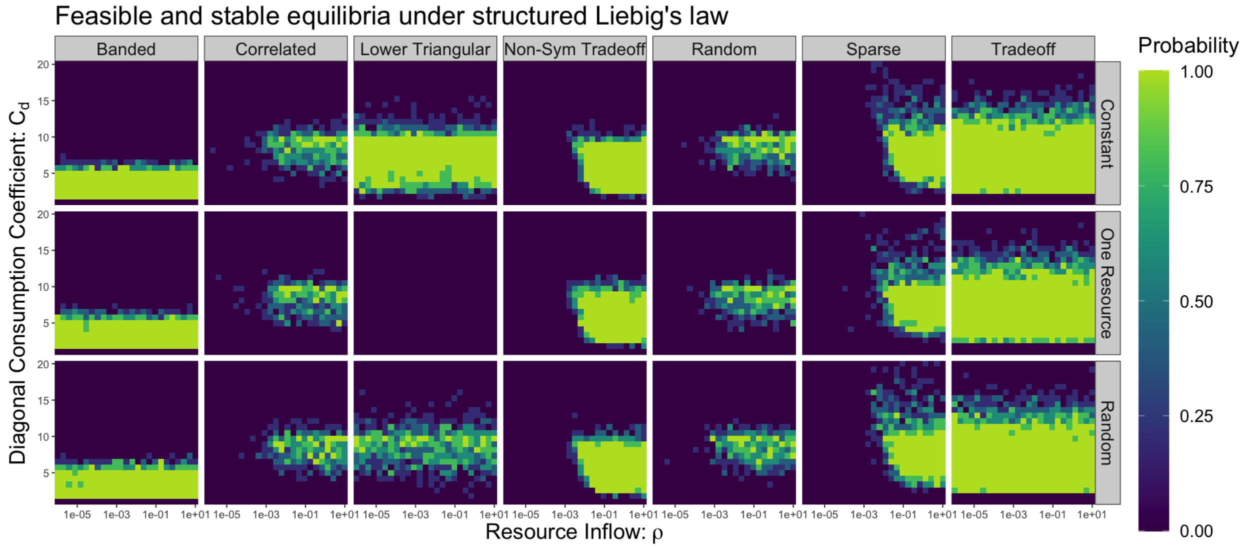

Figure G: **Probability of feasible and stable equilibria under structured Liebig's law.** Parameters and matrix parameterizations are exactly as in Fig. F except now consumer grow according to the structured version of Liebig's law

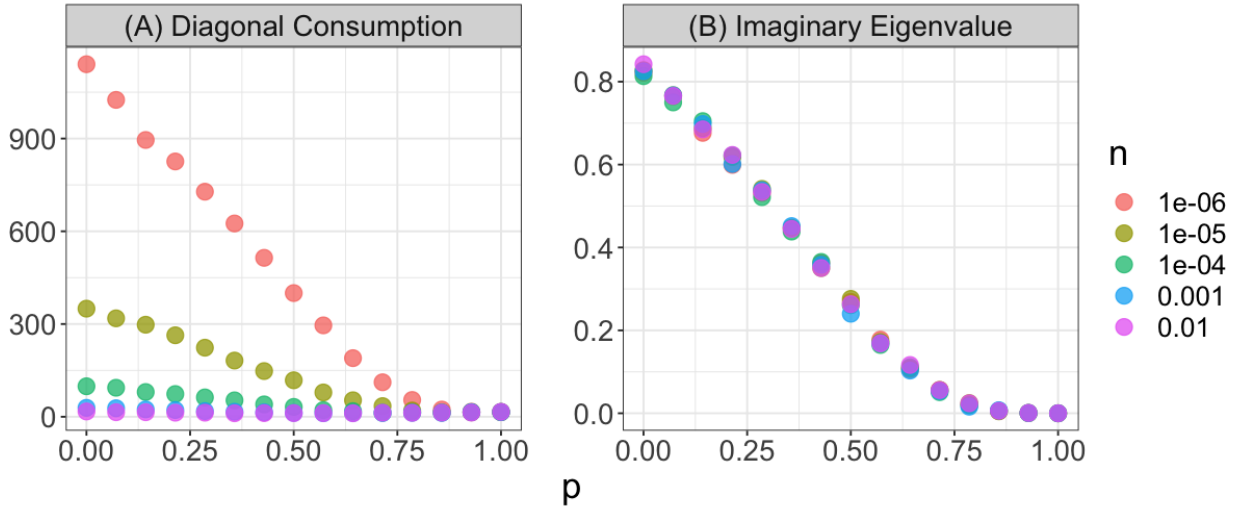

Figure H: **Off-diagonal correlation between the consumption coefficients promotes stability.** (A) We plot the smallest value of $C_d$ at which the system becomes unstable or unfeasible averaged over $5$ replicates as a function of the parameter $p$. In the correlated $C$ matrix case, $p$ determines the correlation between the pair $(C_{ij}, C_{ji})$ by the formula $C_{ji} = pC_{ij} + (1-p)C'_{ij}$ where $C_{ij}$ and $C'_{ji}$ are uniform random variables on $[0, 1]$. We take $P$ as in the constant case. The different colors are different values of the consumer abundance $n$. As the consumer abundances decrease, $C_d$ must be large to maintain stability for uncorrelated matrices (ie. small $p$). However, when $p$ is near $1$, and hence the off-diagonal pairs $(C_{ij}, C_{ji})$ are highly correlated, $n$ no longer affects the numerically determined value of $C_d$. Because the transition between these two regimes is smooth, even approximate symmetry in the $C$ matrix promotes stability. (B) For the same range of $p$ values and choices of consumer abundance $n$, we plot the average of the absolute value of the imaginary part of the eigenvalues of the matrix $B = -C + P^T C$. This quantity does not depend on $n$, but decreases to zero as $p$ goes to $1$ when the matrix $B$ has purely real eigenvalues. Parameters: $S = 15$, $\epsilon = 0.9$ and $r = 1$.

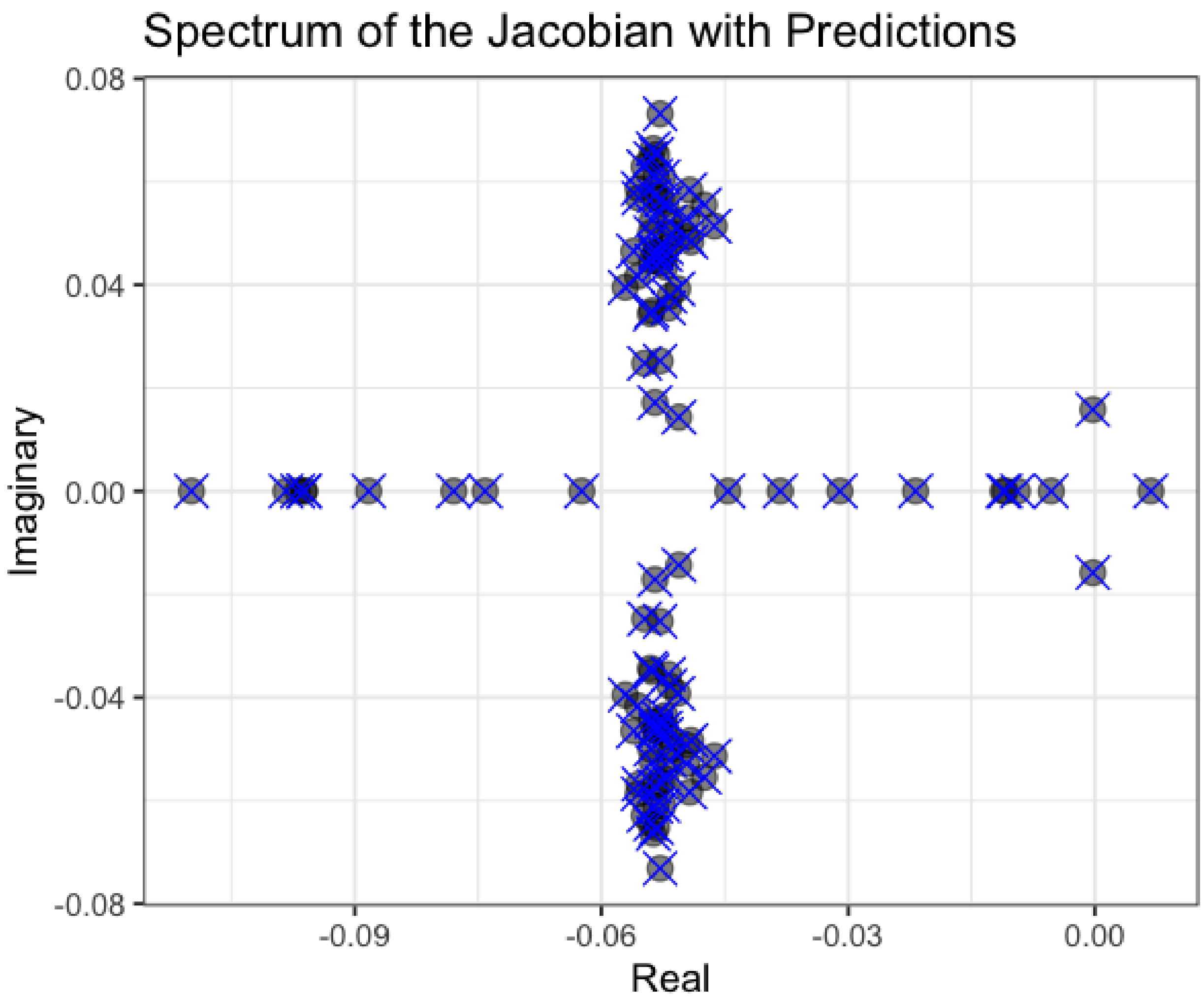

Figure I: **Predicting the spectrum of $J$ from the spectra of $A$ and $B$ when $A$ and $B$ are simulataneously diagonalizable.** Gray circles are the spectrum of $J$ and blue crosses are analytical predictions from the spectra of $A$ and $B$. Parameters: $S = 100$, $C_d = 10$, $\epsilon = 0.05$, $n = 0.001$ and $r = 1$. $C$ and $P$ are given by the circulant parameterization with an underlying uniform distribution on $[0, 2]$ and $[0, 1]$ respectively before the constraints are imposed.

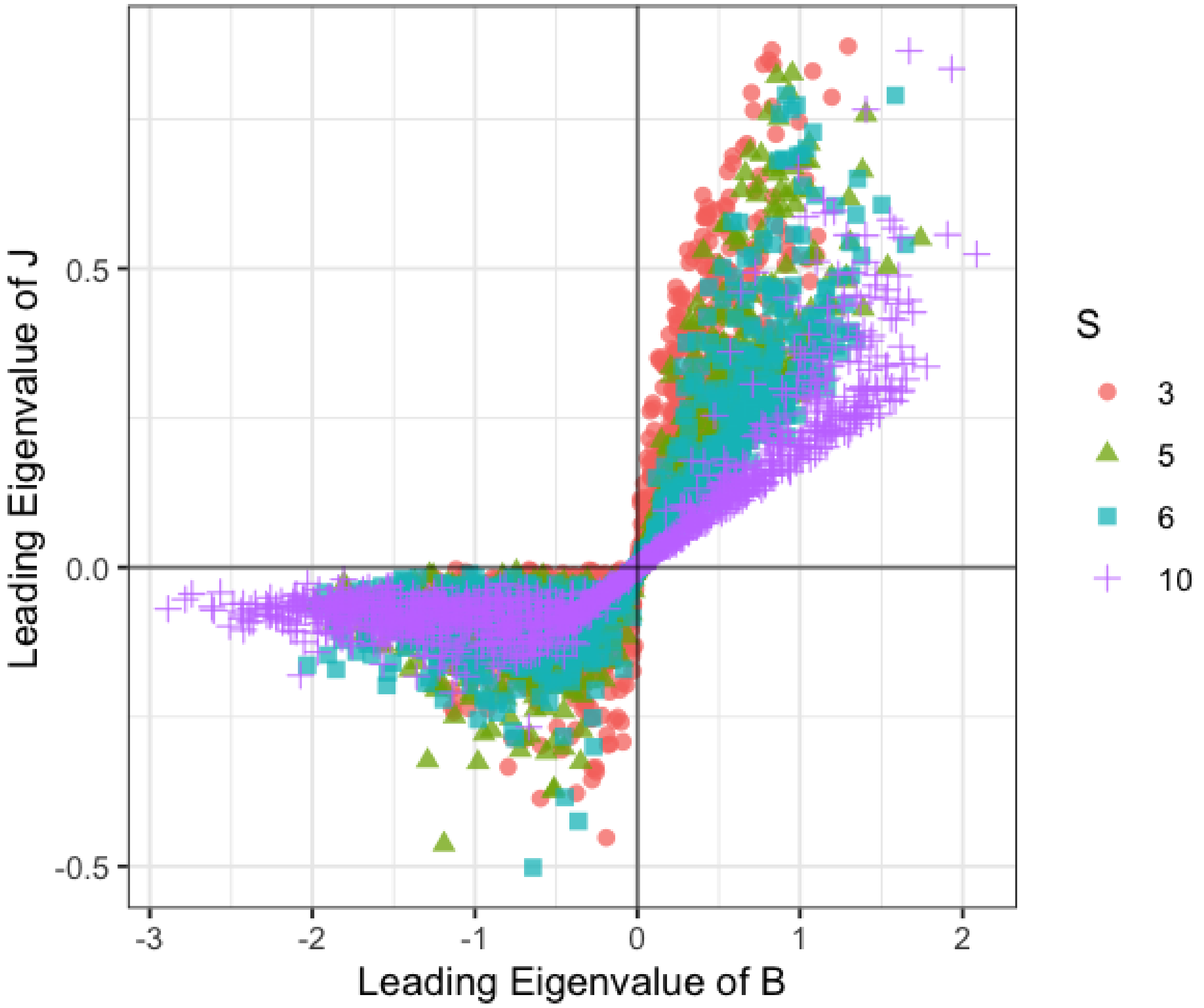

Figure J: $J$ **is stable if and only if** $B$ **is stable in simulations** We plot the eigenvalue of $J$ with maximum real part as a function of the largest eigenvalue of $B$ for $1000$ replicates. The different colors and types of points indicate different number of species $S$. The sign of the leading eigenvalue of $B$ is always the same as the sign of the real part of the leading eigenvalue of $J$, and the transition from stability to instability (negative to positive values) appears to be sharp.

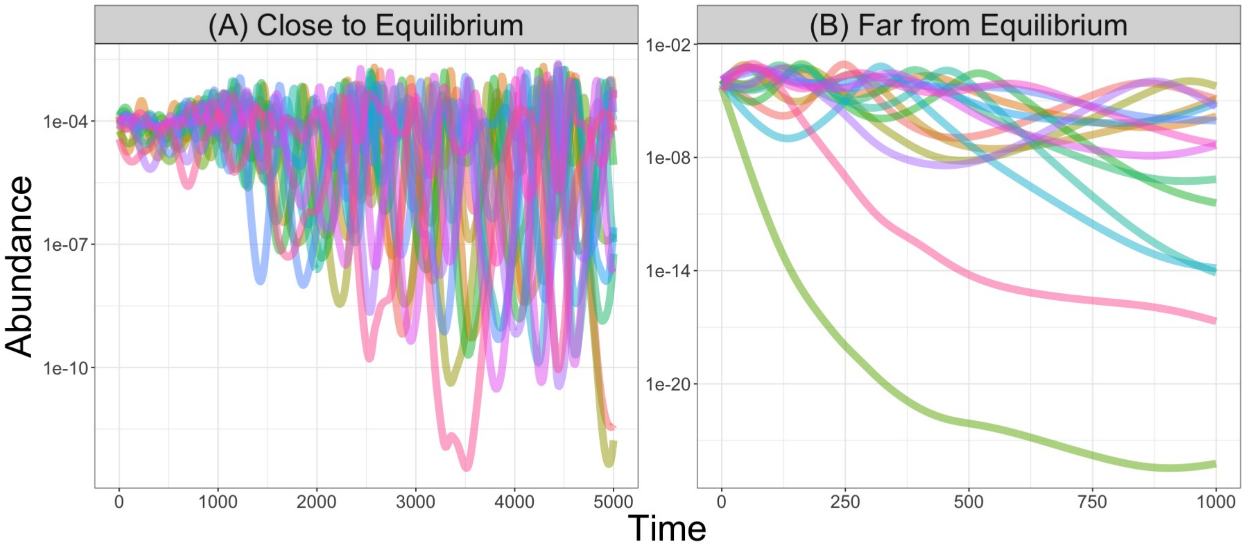

Figure K: **Consumer dynamics close to and far from an unstable equilibrium.** We plot the consumer dynamics over time when the equilibrium is unstable for initial conditions close to (A) and far from (B) equilibrium. To determine the initial conditions, we add noise to the equilibrium values of $\vec{N}$ and $\vec{R}$. We generate $2S$ samples from a normal distribution with mean $0$ and a specified standard deviation ($0.01$ in panel (A) and $0.1$ in panel (B)). We then multiply these random samples by the equilibrium abundances and add this quantity to the equilibrium abundances to get the initial conditions. Parameters: $S = 15$, $C_d = 5$, $\epsilon = 0.05$ and $\vec{\eta} = \vec{1}$. The $C$ matrices are sampled according to the sparse matrix parameterization with an underlying uniform distribution on $[0.5, 1.5]$. The matrix $P$ is given by the constant parameterization. The resource inflow was $\vec{\rho} = 0.0001\vec{1}$. These parameters are the same as those used to generate the unstable dynamics in Fig. 1 of the main text. The stable dynamics in Fig. 1 are also the same parameters except with the total resource inflow $\rho = 1$.

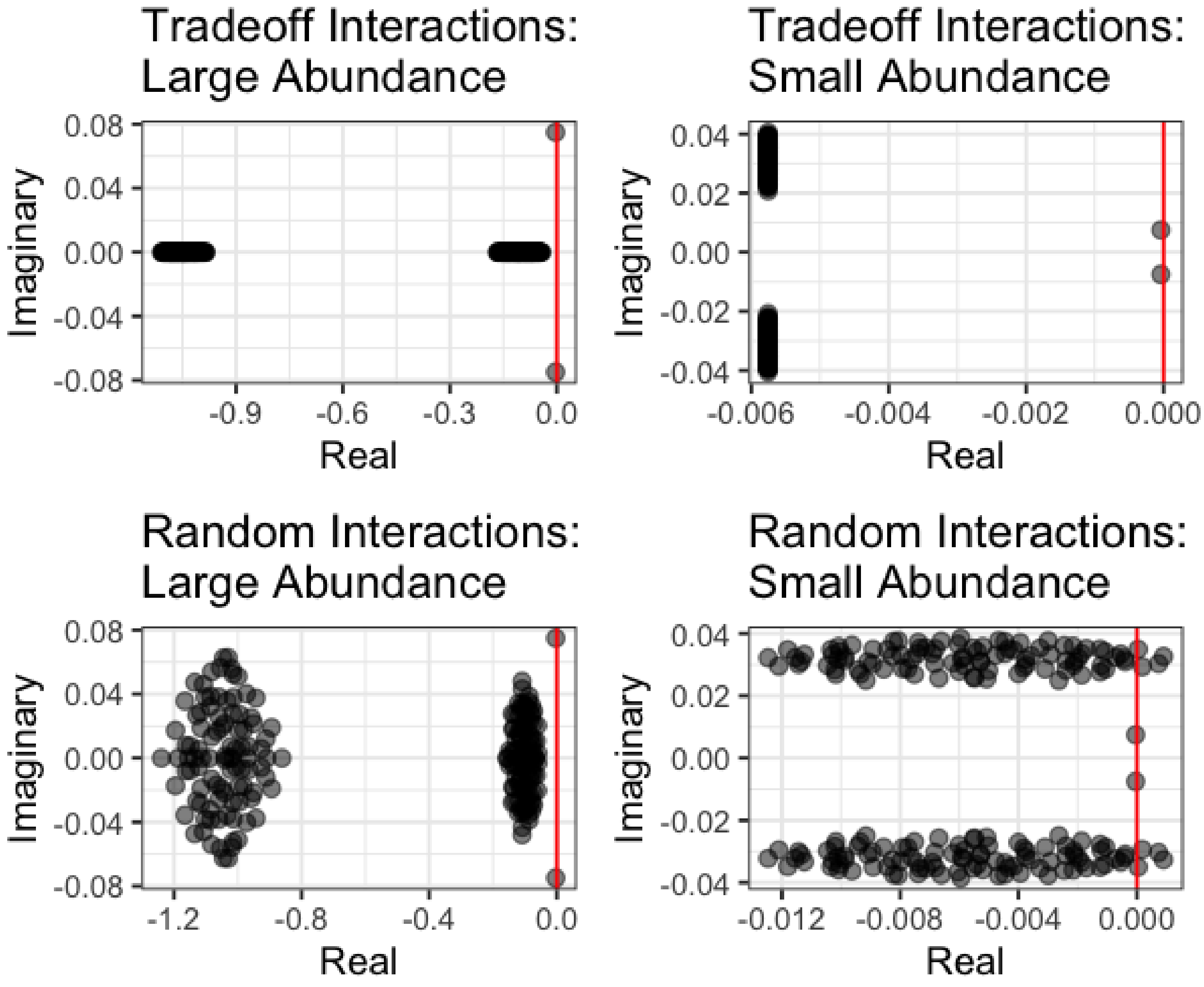

Figure L: **The spectrum of $J$ for small and large consumer abundances.** We plot the spectrum of $J$ when the consumer abundances are small $n = 0.0001$ and large $n = 0.01$ under the tradeoff and random parameterizations. Red vertical lines indicate the imaginary axis. In all four cases, there are two bulks of eigenvalues, which are each a transformed version of the spectrum of $B$. When the consumer abundances are small, these two eigenvalue bulks appear as two clouds of eigenvalues with non-zero imaginary parts for both parameterizations. For the tradeoff parameterization, the clouds have no width, because the underlying $B$ matrix has purely real eigenvalues. For the random parameterization, the clouds have non-zero width because $B$ has eigenvalues with non-zero imaginary part. As the consumer abundances become small, the non-zero width of these clouds means that an eigenvalue complex conjugate pair eventually crosses the imaginary axis at a value $n > 0$. Parameters: $S = 100$, $C_d = 15$, $\epsilon = 0.05$ and $r = 1$. In each of the parameterizations, the consumption coefficients are sampled from a uniform distribution on $[0, 2]$ before the constraints are imposed. In the tradeoff parameterization, the production matrix is constant, while in the random parameterization, the production coefficients are sampled from a uniform distribution on $[0, 1]$ before the constraints are imposed.
